# Distinct and compensatory roles of Stag1 and Stag2 in post-mitotic genome refolding

**DOI:** 10.1101/2025.10.24.684407

**Authors:** Manzhu Wang, Fengnian Shan, Lirong Shu, Sijian Xia, Fuhai Liu, Dannan Jing, Yabing Zhang, Yongjia Weng, Bicheng Li, Han Zhao, Yinzhi Lin, Yali Yu, Baiyue Wang, Haoyue Zhang

## Abstract

The three-dimensional architecture of the eukaryotic genome is largely shaped by the cohesin complex, which contains either Stag1 or Stag2 subunits. Although both subunits contribute to chromatin organization, their specific functions in *de novo* loop formation during post-mitotic genome refolding remain elusive. Here, we leverage the mitosis-to-G1 transition to dissect their individual roles. We found that Stag1 depletion has a negligible impact on post-mitotic genome restructuring or transcription reactivation. In contrast, Stag2 orchestrates chromatin remodeling in a manner that is both cell cycle stage-specific and chromatin context-dependent. During early-G1, Stag2 preferentially associates with euchromatin, where it drives the rapid formation of small chromatin loops. This facilitates prompt promoter-enhancer (P-E) contact formation and enables efficient transcription activation. As the nuclear concentration of Stag2 increases by late-G1, it progressively suppresses large loops, likely due to its shorter chromatin residence time and its potential to competitively displace the more extrusion-capable Stag1-associated cohesin. Mechanistically, Stag2-mediated loop extrusion is constrained by CTCF-bound barriers, rather than by genomic travel distance. Although Stag2 associates rapidly with euchromatin in early-G1, its recruitment to heterochromatin is delayed until late-G1. Simultaneous depletion of both Stag proteins results in a synergistic loss of virtually all structural loops and a more severe disruption of transcription than that caused by individual deletions. Together, these results establish Stag2 as the principal regulator of post-mitotic genome reorganization among Stag paralogs, mediating spatiotemporal control of chromatin architecture, while Stag1 provides compensatory support to ensure functional robustness.

## Introduction

The eukaryotic genome is intricately folded, exhibiting a multi-layered hierarchical three-dimensional (3D) structure. At the megabase scale, chromatin can be partitioned into the euchromatin-enriched A-type compartments and heterochromatin-enriched B-type compartments^1-3^. At finer scales, chromatin is characterized by topologically associating domains (TADs) and chromatin loops^1,4-6^. One major organizer of interphase genome structure is cohesin, a multi-subunit complex initially known for its role in cohering sister chromatids after DNA replication ^6,7^. Recent work has revealed another critical role of cohesin in interphase: organizing genome architecture via a process termed loop extrusion. In this process, cohesin actively translocates along the DNA fiber, forming progressively larger loops until it is arrested by convergently oriented CTCF molecules^8-11^. By maintaining these structures, cohesin and CTCF cooperate to either facilitate or insulate contacts between gene promoters and distal enhancers, thereby regulating transcription^6,12-14^.

The cohesin complex consists of two structural maintenance of chromosome proteins (SMC1 and SMC3), a kleisin subunit (Rad21), and one of the two paralogous Stag subunits, Stag1 or Stag2^15,16^. Despite sharing broadly overlapping functions, accumulating evidence indicates that Stag1- and Stag2-associated cohesin exhibit specialized roles in genome organization and transcription regulation^17,18^. For instance, while both Stag proteins occupy a common set of binding sites on chromatin, they also display unique genomic positioning^19^. Furthermore, Stag1-associated cohesin exhibits significantly longer chromatin residence time than Stag2-associated cohesin^20^. Therefore, Stag1 may be primarily involved in establishing stable, long-range chromatin loops. In contrast, Stag2-associated cohesin may mediate transient, short-range interactions, including those between enhancers and promoters within TADs^19,20^. Consistent with these findings, deleting Stag2 results in enlarged chromatin loops, likely due to the increased availability of Stag1-associated cohesin, which favors large loop formation^20^. Despite these discoveries, how each Stag protein contributes to the *de novo* establishment of structural loops remains unknown.

3D genome structure is dynamically remodeled throughout the cell cycle. During mitosis, hallmark chromatin architectural features including A/B compartments, TADs and chromatin loops are disrupted^21^. During this process, cohesin is evicted from chromatin^22,23^. As cells exit from mitosis, interphase genome structure is gradually rebuilt, accompanied by a progressive reloading of cohesin onto the genome, which drives chromatin re-looping^24^. A recent study utilizing a quantitative imaging approach illustrates that Stag1 is rapidly imported into daughter nuclei and rebinds chromatin, while Stag2 nuclear import is delayed, leading to a gradual chromatin binding over a course of several hours into G1 phase^25^. This temporal uncoupling may indicate stage-specific roles of Stag1 and Stag2 in post-mitotic genome refolding.

Here, we adopt the mitosis-to-G1 transition as a platform to systematically dissect how Stag1- and Stag2-associated cohesin independently regulate chromatin structural loop reformation and gene reactivation after cell division. We find that Stag1 and Stag2 share redundant and unique functions. Stag1 depletion elicits no dramatic effect on post-mitotic chromatin loops or transcription. Conversely, Stag2 exhibits two regulatory modes: (1) a loop size-dependent regulation, in which it constitutively stabilizes small loops (<100 kb) throughout G1 phase while progressively suppressing large loops (>300 kb) as its nuclear concentration increases; and (2) a chromatin context-dependent regulation, in which it rapidly engages euchromatin in early-G1 to ensure efficient transcription reactivation, followed by a delayed association with heterochromatin in late-G1. Co-depletion of both Stag proteins triggered synergistic collapse of structural loops, revealing a functional compensation between them. These results delineate Stag2 as the primary architect among Stag paralogs regulating post-mitotic genome refolding.

## Results

### Impacts of Stag1 and Stag2 on post-mitotic chromatin re-compartmentalization

To examine the roles of Stag1 and Stag2 in the re-establishment of chromatin structure following mitosis, we aimed to selectively delete them during the transition from mitosis to G1-phase. To achieve this, we utilized CRISPR/Cas9-directed genome editing on the well-characterized G1E-ER4 erythroblast line^26,27,28^. Two sublines were generated, each carrying a minimal auxin-inducible degron (mAID) endogenously fused to either Stag1 (*Stag1*^mAID^) or Stag2 (*Stag2*^mAID^) (Fig. 1a)^29,30^. 5-Ph-IAA treatment led to complete degradation of the target proteins within 4 hours as assessed by western blot analysis and chromatin immunoprecipitation and sequencing (ChIP-seq) (Extended Data Fig. 1a-c). Notably, prolonged depletion of Stag1 did not delay cell proliferation (Extended Data Fig. 1d). In contrast, Stag2 loss led to a markedly slower cell growth (Extended Data Fig. 1d). Intriguingly, Stag1 and Stag2 depletion each left the other’s chromatin-binding profile unchanged, suggesting that their genomic occupancies are independent of each other (Extended Data Fig. 2a-c).

**Figure 1:**
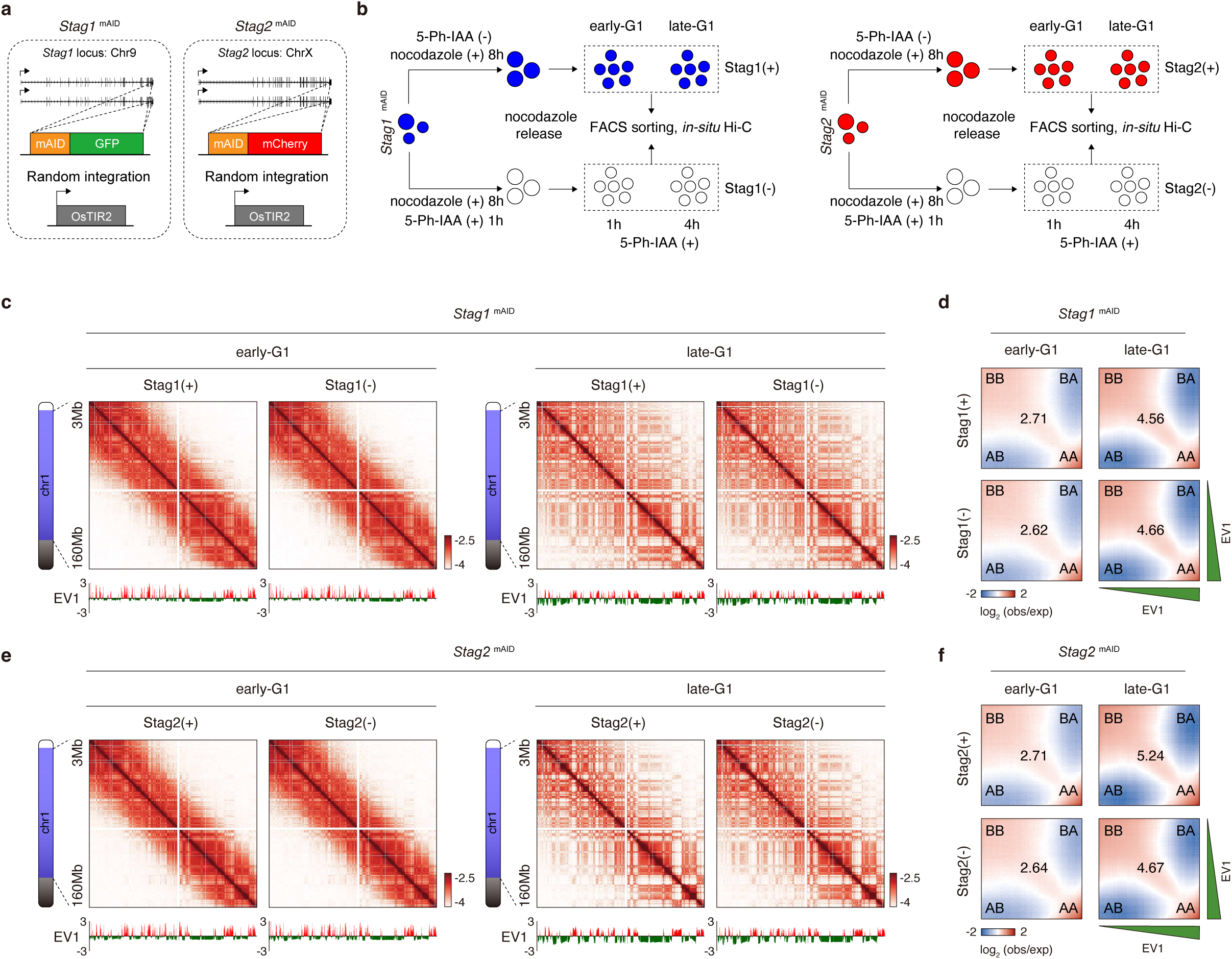
Post-mitotic chromatin re-compartmentalization in the absence of Stag1 or Stag2. **a**, Schematic of *Stag1*^mAID^ and *Stag2*^mAID^ cell line construction. **b,** Workflow for mitotic arrest/release and 5-Ph-IAA treatment to obtain early-G1 and late-G1 populations with or without Stag1 or Stag2. **c**, KR-balanced, 100 kb-binned Hi-C contact maps of showing post-mitotic chromatin re-compartmentalization with or without Stag1. EV1 tracks of corresponding contact maps are shown in parallel. **d**, Saddle plots showing the comparable compartment scores before and after Stag1 depletion. **e**, KR-balanced, 100kb-binned Hi-C contact maps showing post-mitotic chromatin re-compartmentalization with or without Stag2. EV1 tracks are shown below each contact map. **f**, Saddle plots showing slightly reduced compartment score in late-G1 phase upon Stag2 depletion.

The acute nature of mAID-mediated protein degradation enabled us to investigate the effects of Stag1 or Stag2 depletion during cell division. *Stag1*^mAID^ or *Stag2*^mAID^ cells were arrested at prometaphase through nocodazole treatment (Fig. 1b). Meanwhile, 5-Ph-IAA was applied to allow protein degradation (Fig. 1b). Upon nocodazole release, cells devoid of Stag1 or Stag2 were collected at 1h (early-G1) and 4h (late-G1) time points, respectively, using fluorescence-activated cell sorting (FACS) (Fig. 1b). Cells without 5-Ph-IAA treatment were collected in parallel as controls (Fig. 1b). Of note, short-term depletion of Stag proteins did not affect mitotic progression (Extended Data Fig. 1e).

It is well-established that cohesin-mediated loop extrusion counteracts chromatin compartmentalization^31,32^. To test whether deleting Stag proteins could affect post-mitotic chromatin re-compartmentalization, we performed *in-situ* Hi-C experiments. *In-situ* Hi-C on average yielded ∼200 million valid interaction pairs for each sample with high concordance among biological replicates (Extended Data Fig. 3a; Supplementary Table 1). Consistent with prior reports, the control samples exhibited the characteristic checkerboard-like A/B compartments in early-G1 phase, which were progressively intensified and expanded across the entire genome by late-G1 phase (Fig. 1c, e)^24,33^. Deleting Stag1 did not induce any noticeable changes in post-mitotic chromatin re-compartmentalization (Fig. 1c, d; Extended Data Fig. 3b-d, Extended Data Fig. 4a). Similarly, Stag2 depletion had a minimal impact on the overall compartmentalization of the genome following mitosis (Fig. 1e, f; Extended Data Fig. 3c). However, we observed a mild reduction of A/B compartment segregation specifically during late-G1 upon Stag2 depletion (Extended Data Fig. 4b, c), which was not seen in Stag1-deficient cells (Extended Data Fig. 4a). This finding contrasts with previous reports that complete cohesin loss (e.g., via Rad21 depletion) sharpens the separation between A and B compartments^31,34^. We reason that the remaining Stag1-associated cohesin, which possesses longer chromatin residence time, not only compensates for the loss of Stag2-mediated loop extrusion but does so with enhanced activity, resulting in a net decrease in A/B segregation. Together, our data unveil distinct roles of Stag1 and Stag2 in regulating higher-order chromatin re-organization after mitosis.

### Stag1 depletion slightly attenuates large loop reformation after mitosis

Reformation of structural loops after mitosis is tightly controlled by cohesin recruitment ^24,33,35^. Although cohesin reloading is generally a gradual process post-mitosis, Stag1- and Stag2-associated complexes exhibit strikingly different dynamics: Stag1 reloads rapidly onto chromatin, while Stag2 displays delayed nuclear import and more gradual recruitment^25^. Yet, whether these divergent reloading kinetics translate into distinct functional roles of Stag1 and Stag2 in directing post-mitotic chromatin re-looping remains unclear. To address this, we employed a modified HICCUPS approach to systematically identify chromatin loops in early- and late-G1 phase samples, both with and without Stag proteins. In total, we identified 23,482 non-redundant chromatin loops with high concordance among biological replicates (Extended Data Fig. 5a). Among these, 9,428 were classified as structural loops, characterized by co-occupancy of CTCF and cohesin peaks at both anchors with at most one anchor marked by cis-regulatory elements (CREs). Surprisingly, aggregated peak analysis (APA) of genome-wide structural loops revealed that Stag1 depletion did not measurably alter post-mitotic structural loop strength (Extended Data Fig. 5b). In line with these results, the distance-dependent contact probability decay curve (*P_S_*) was unchanged upon Stag1 depletion after mitosis, suggesting that overall cohesin loop extrusion activity was unaffected (Extended Data Fig. 5c). Intriguingly, closer examination uncovered a subtle reduction of loop strength, particularly for large loops (Extended Data Fig. 5d-f). This trend was observed in both early- and late-G1 cells (Extended Data Fig. 5d-f). The early-G1 effect was consistent with the rapid post-mitotic recruitment of Stag1^20,25^. Together, these results indicate that Stag1 slightly contributes to post-mitotic genome refolding, with a specific role in the efficient formation of large loops.

### Stag2 plays a bifurcated role in post-mitotic loop reformation

The mild impact of Stag1 loss on post-mitotic loop reformation implies a functional compensation by Stag2. This aligns with the status of Stag2 as the dominant paralog, accounting for ∼75% of the total cohesin pool^36^. To investigate Stag2’s specific contributions, we measured the dynamic reformation of chromatin loops in Stag2-deficient cells after mitosis. Our analysis revealed that 1,624 structural loops were gained upon Stag2 depletion (Fig. 2a; Extended Data Fig. 6a). Of note, the gained loops were detected almost exclusively in late-G1 phase and were absent in early-G1 (Fig. 2b). This delayed phenotype is consistent with the gradual nuclear accumulation of Stag2-associated cohesin. Furthermore, the gained structural loops were significantly larger than those that were maintained or lost, confirming previous findings that Stag2 depletion promotes large loop formation (Fig. 2c)^20^. Conversely, 973 loops were lost following Stag2 depletion (Fig. 2a; Extended Data Fig. 6a). These lost loops were significantly smaller than both maintained and gained ones (Fig. 2c). This implies that Stag2 plays a role in establishing small structural loops after mitosis, a function distinct from its suppressive effect on large loop formation. To further elucidate the impact of Stag2 loss on loops of different sizes, we systematically quantified the log_2_ fold change (log_2_FC) of loop intensity across all identified structural loops. Our analysis revealed that Stag2 depletion triggers two distinct, loop size-dependent effects that are temporally uncoupled during G1 phase. First, large loops (>300 kb) were predominantly intensified by Stag2 loss in late-but not early-G1 phase (Fig. 2d, e; Extended Data Fig. 6b). Second, small loops (<100 kb) were attenuated rapidly in early-G1 upon Stag2 degradation and remained weakened throughout G1 phase (Fig. 2d, e; Extended Data Fig. 6b).

**Figure 2:**
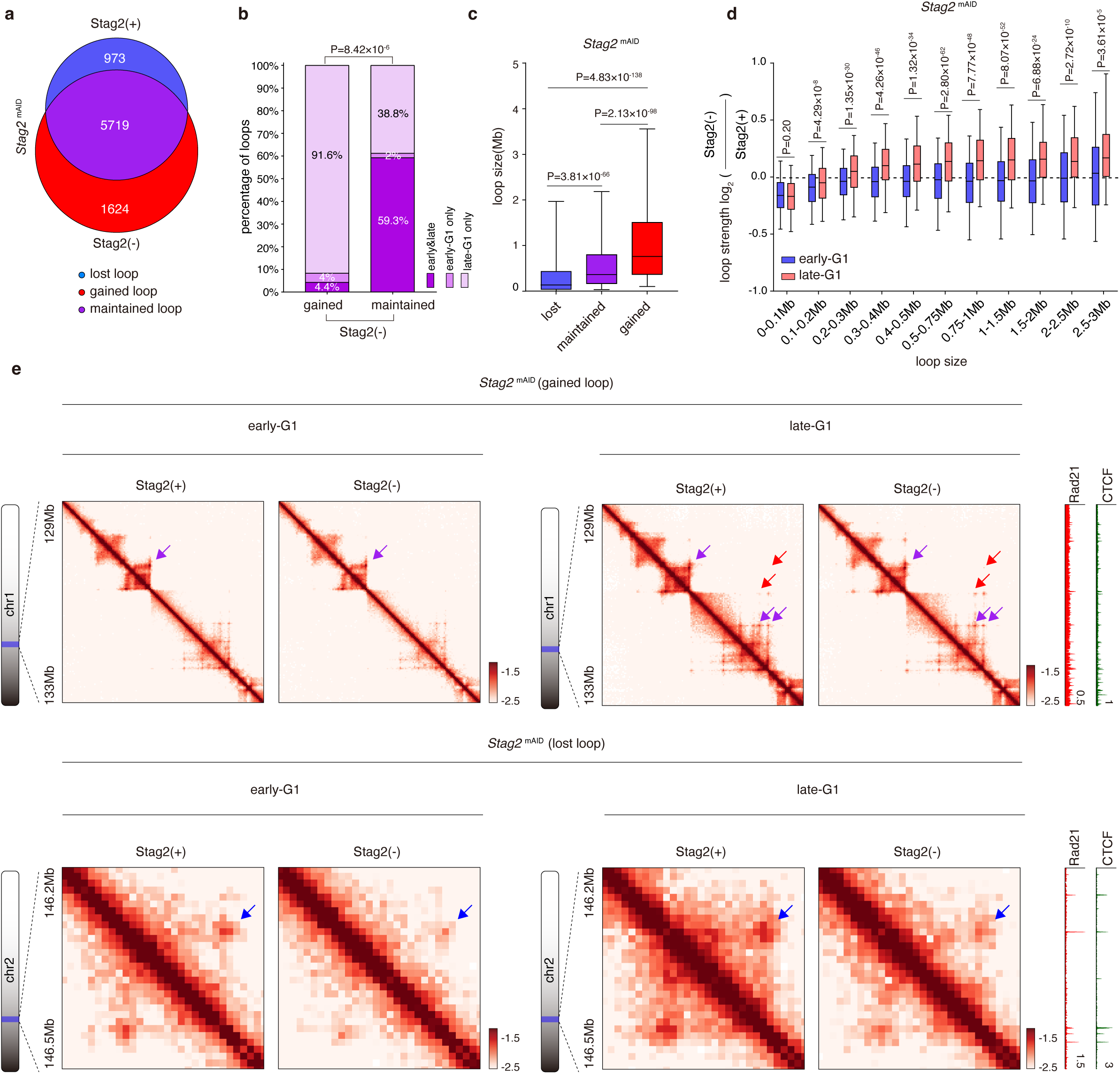
Stag2 depletion alters post-mitotic structural loop reformation. **a**, Venn diagram showing the number of lost, maintained and gained structural loops upon Stag2 depletion after mitosis. **b**, Bar graphs showing the percentage of gained and maintained loops classified by their presence in early-G1, late-G1, or both stages. *P* value is calculated using a two-sided Fisher’s exact test. **c**, Boxplots showing size distributions of the lost, maintained, and gained structural loops upon Stag2 depletion. *P* values were calculated using two-sided Wilcoxon rank-sum test. **d**, Boxplots showing the log_2_ fold change (FC) of the strength of structural loops with indicated size-ranges upon Stag2 depletion. The log_2_FC for both early- and late-G1 phase were shown. *P* values were calculated using two-sided paired Wilcoxon rank-sum test, comparing early-vs late-G1 log_2_FC for the same loops within each size bin. **e**, KR-balanced, 10kb-binned Hi-C contact maps showing representative gained loops and lost loops in early-G1 and late-G1 phase. Lost, maintained and gained loops are indicated by blue, purple and red arrows respectively. Late-G1 phase CTCF and Rad21 ChIP-seq tracks shown on the right.

Our findings support a model in which Stag2 orchestrates post-mitotic chromatin re-looping through a dual mechanism. During early-G1, its low nuclear abundance allows the establishment of small loops. As Stag2 progressively accumulates by late-G1, this increased concentration allows it to both maintain small loops and competitively displace Stag1-associated cohesin, thereby suppressing large loop formation. Thus, the kinetics of Stag2 nuclear import could dictate the temporal partitioning of its functions in 3D genome re-organization after mitosis.

### Intermediate CTCF roadblocks but not genomic distance constrains Stag2-mediated loop formation

We propose that the Stag2-associated cohesin complex is intrinsically limited in its ability to form large structural loops, owing to its shorter chromatin residence time relative to Stag1-cohesin. The formation of large chromatin loops is inherently challenged by (1) the need to traverse long genomic distances without dissociating from chromatin and (2) the need to bypass multiple intermediate CTCF sites that may prematurely arrest extrusion (Fig. 3a).

**Figure 3:**
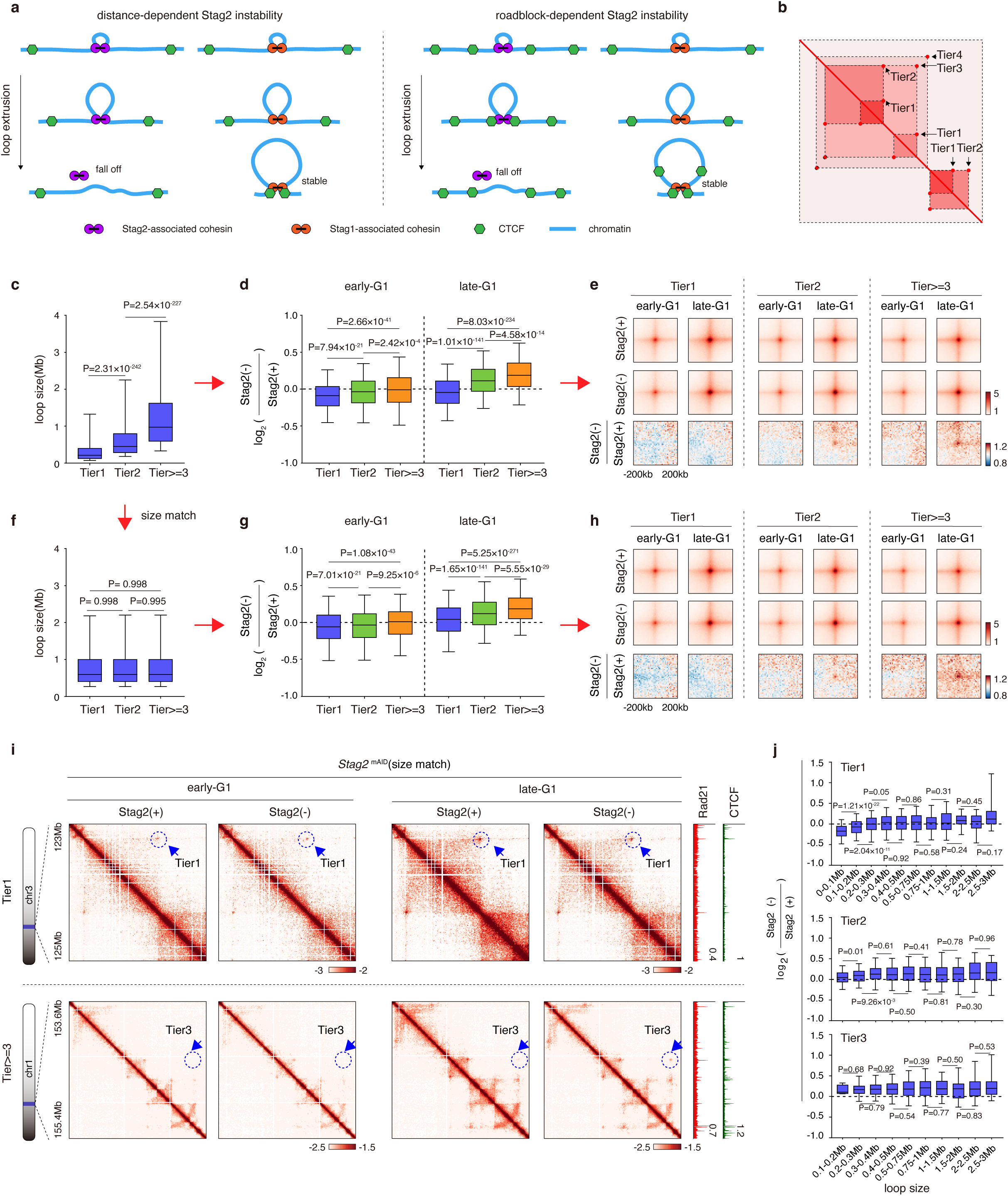
Intermediate CTCF roadblocks undermine Stag2-mediated large loop formation. **a**, Schematic illustration showing the two potential factors constraining Stag2’s capacity to form large loops. **b**, Schematic illustration showing the hierarchical organization of structural loops (Tier1, Tier2, and Tier≥3). **c**, Boxplots showing the size distributions of Tier1, Tier2, and Tier≥3 structural loops. **d**, Boxplots showing the effects of Stag2 depletion on the strength of structural loops of indicated tiers at early- and late-G1 phase. *P* values were calculated using two-sided Wilcoxon rank-sum test. **e**, Aggregate peak analysis (APA) plots showing the changes of structural loops of indicated tiers upon Stag2 loss in early- and late-G1 phase. **f**–**h**, Similar to (**c**–**e)**, showing the same trend of structural loop change after size-matching. **i**, Representative Hi-C contact maps of a pair of size-matched Tier1 and Tier3 loops early-G1 and late-G1 before and after Stag2 depletion. Late-G1 CTCF and Rad21 ChIP-seq tracks are shown on the right. **j,** Boxplots showing the log_2_FC in structural loop strength upon Stag2 depletion in late-G1 phase. Three separate plots, each showing one tier are presented, with loops organized by size-ranges. *P* values were calculated using two-sided Wilcoxon rank-sum tests.

To dissect the biophysical constraints limiting Stag2-associated cohesin’s loop forming capacity, we employed a hierarchical analysis and stratified structural loops into three hierarchical categories: Tier1 (primary, nested loops), Tier2 (loops encompassing Tier 1), and Tier≥3 (higher-order loops), positing that higher-tier loops present more CTCF-mediated obstacles (Fig. 3b). We reasoned that if Stag2 is inherently restricted in forming large loops, then its depletion should strengthen such loops by increasing the pool of the more extrusion-capable Stag1-associated cohesin. Stag2 depletion in late-G1 resulted in strengthened Tier2 and Tier≥3 loops, suggesting a limited capacity of Stag2 to form these structures (Fig. 3c–e; Extended Data Fig. 7a– c). It is noteworthy that higher-order loops are usually larger than lower-order ones (Fig. 3c). To specifically isolate the contribution of CTCF roadblocks independent of loop length, we performed a size-matched analysis by randomly selecting loops within the same size range across all three hierarchical groups (Fig. 3f). Despite equivalent sizes, higher-order loops still showed significantly greater enhancement upon Stag2 depletion compared to nested ones (Fig. 3g-i; Extended Data Fig. 7d). This demonstrates that intermediate CTCF roadblocks could undermine Stag2’s loop-forming capacity.

We next investigated whether genomic travel distance independently limits Stag2-cohesin. To isolate distance effects from CTCF roadblock interference, we analyzed loops within each hierarchical tier separately. With Tier1 loops, which are largely devoid of nested obstacles, Stag2 depletion preferentially reduced the intensity of smaller loops, consistent with its established role in promoting small loop formation (Fig. 3j). Interestingly, longer Tier1 loops (>300kb) remained largely unaffected (Fig. 3j). This finding stands in stark contrast to the pronounced enhancement of large loops observed in unstratified loop population, underscoring the necessity of controlling for hierarchical context when interpreting size-dependent extrusion phenotypes (Fig. 2d). Within Tier2, a modest distance-dependent increase in intensity was detected for loops between 100 to 400 kb upon Stag2 loss (Fig. 3j). However, this sensitivity was completely abolished in Tier3 loops (Fig. 3j). Collectively, these findings indicate that while travel distance can modestly affect Stag2-mediated loop formation in the absence of intermediate barriers, this effect is overridden by the presence of CTCF roadblocks, which constitute the dominant constraint on Stag2’s loop-forming capacity.

### Chromatin context-dependent regulation of loop extrusion by Stag2

We next asked whether loop extrusion by Stag2-associated cohesin is influenced by the epigenetic context of chromatin. To isolate chromatin context-specific effects from the confounding influence of CTCF roadblocks, we focused our analysis on Tier1 loops. We classified these loops into three distinct epigenetic categories using unsupervised *k-means* clustering: the H3K27ac and H3K36me3 enriched euchromatin loops (cluster1, n=1207), the H3K27me3 enriched facultative heterochromatin loops (cluster2, n=1545), and the H3K9me3 enriched constitutive heterochromatin loops (cluster3, n=782) (Fig. 4a). For each type of structural loops, we evaluated two distinct readouts: (1) puncta signals, representing the relatively stable, CTCF-anchored loop structures, and (2) internal contacts within loop bodies, reflecting dynamic, transient extruding intermediates mediated by cohesin (Fig. 4b).

**Figure 4:**
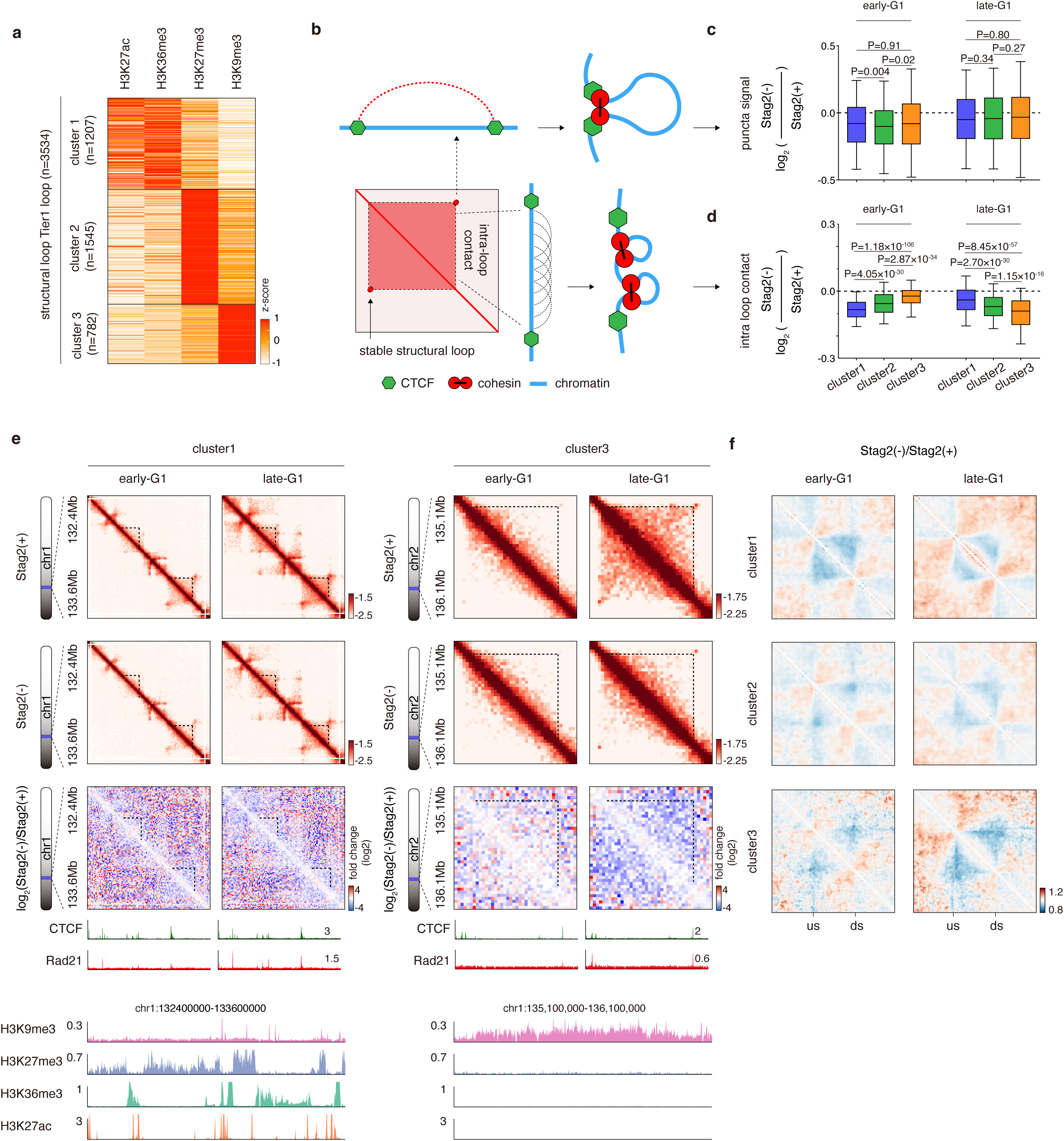
Chromatin context-dependent regulation of loop extrusion by Stag2. **a**, Heatmap showing the clustering result of Tier 1 structural loops based on indicated histone modifications. **b**, Schematic illustration showing the representative chromatin conformation, constituting the puncta and intra-loop contact signals in Hi-C contact maps. **c**, Boxplots showing the log_2_FC in puncta signal upon Stag2 depletion across loop clusters. For each phase, *P* values were obtained from two-sided Wilcoxon rank-sum tests for pairwise comparisons between clusters. **d**, Boxplots showing the log_2_FC in intra-loop contacts across loop clusters in early- and late-G1. *P* values were obtained from two-sided Wilcoxon rank-sum tests. **e**, Representative Hi-C contact maps for cluster1 and cluster3 structural loops in early- and late-G1 phase with or without Stag2. Contact maps are accompanied by corresponding CTCF and Rad21 ChIP-seq tracks. Tracks of histone modifications corresponding to the same regions of cluster1 and 3 loops were also shown. Dotted lines indicate intra-loop contacts. **f**, Compiled and rescaled contact maps showing the distinct impact of Stag2 loss on the intra-loop contacts across all three clusters. Contact maps for both early- and late-G1 phase cells are shown. “us” and “ds” denotes the upstream and downstream loop anchors, respectively.

As expected, Stag2 depletion reduced Tier1 loop puncta signals in early-G1, consistent with its role in promoting small loop establishment after mitosis (Fig. 4c). This reduction persisted throughout G1 phase (Fig. 4c). Notably, the decrease was relatively uniform across all three epigenetic loop clusters, with no dramatic differences among them (Fig. 4c). These findings imply a chromatin context-independent role of Stag2 in the establishment and maintenance of stable structural loops after mitosis.

In contrast, internal loop contacts exhibited pronounced spatiotemporal and chromatin context-dependent regulation by Stag2. In early-G1 phase, Stag2 depletion reduced these contacts across all structural loop clusters (Fig. 4d). However, the magnitude of this effect varied dramatically: being the greatest in the euchromatic loops (cluster 1) and mildest in the constitutive heterochromatic loops (cluster 3) (Fig. 4d-f; Extended Data Fig. 8). This observation suggests a preferential role of Stag2 in initiating extrusion in open chromatin. Strikingly, this relationship was completely reversed in late-G1. Upon Stag2 loss, constitutive heterochromatic loops (cluster 3) showed the most severe disruption in intra-loop contacts, while euchromatic loops (cluster 1) were less affected (Fig. 4d–f; Extended Data Fig. 8). This temporal inversion highlights that Stag2 is increasingly required for loop extrusion in repressive chromatin as cells progress into late-G1, likely reflecting its delayed recruitment to these regions. Our study reveals that Stag2 contributes to the formation of stable, CTCF-anchored structural loops across various epigenetic states. Besides, it also mediates dynamic loop extrusion in a spatiotemporally-regulated and chromatin context-specific manner, driving early extrusion in euchromatin and facilitating late extrusion in heterochromatin after mitosis.

### Co-depletion of Stag1 and Stag2 abolishes post-mitotic structural loop reformation

Deleting Stag1 or Stag2 individually exerted limited effects on chromatin loop reformation, suggesting a degree of functional redundancy between them. However, the combined impact of co-depleting both Stag proteins remained largely unexplored. To address this, we engineered a G1E-ER4 subline, in which both Stag proteins were simultaneously tagged with mAID (Fig. 5a). Western blot and ChIP-seq experiments confirmed efficient protein degradation within 4 hours of 5-Ph-IAA treatment (Extended Data Fig. 1a-c). Using the established nocodazole arrest/release protocol, we assessed potential genome folding defects in cells lacking both Stag proteins (Fig. 5b). Notably, unlike the depletion of core cohesin subunits (e.g., Rad21 or SMC1/3), which disrupts sister-chromatid cohesion and blocks cell division, acute loss of Stag1 and Stag2 did not impede mitotic progression (Extended Data Fig. 1e). However, prolonged depletion of both Stag proteins caused a complete cease of cell proliferation, indicating long-term viability defects (Extended Data Fig. 1d).

**Figure 5:**
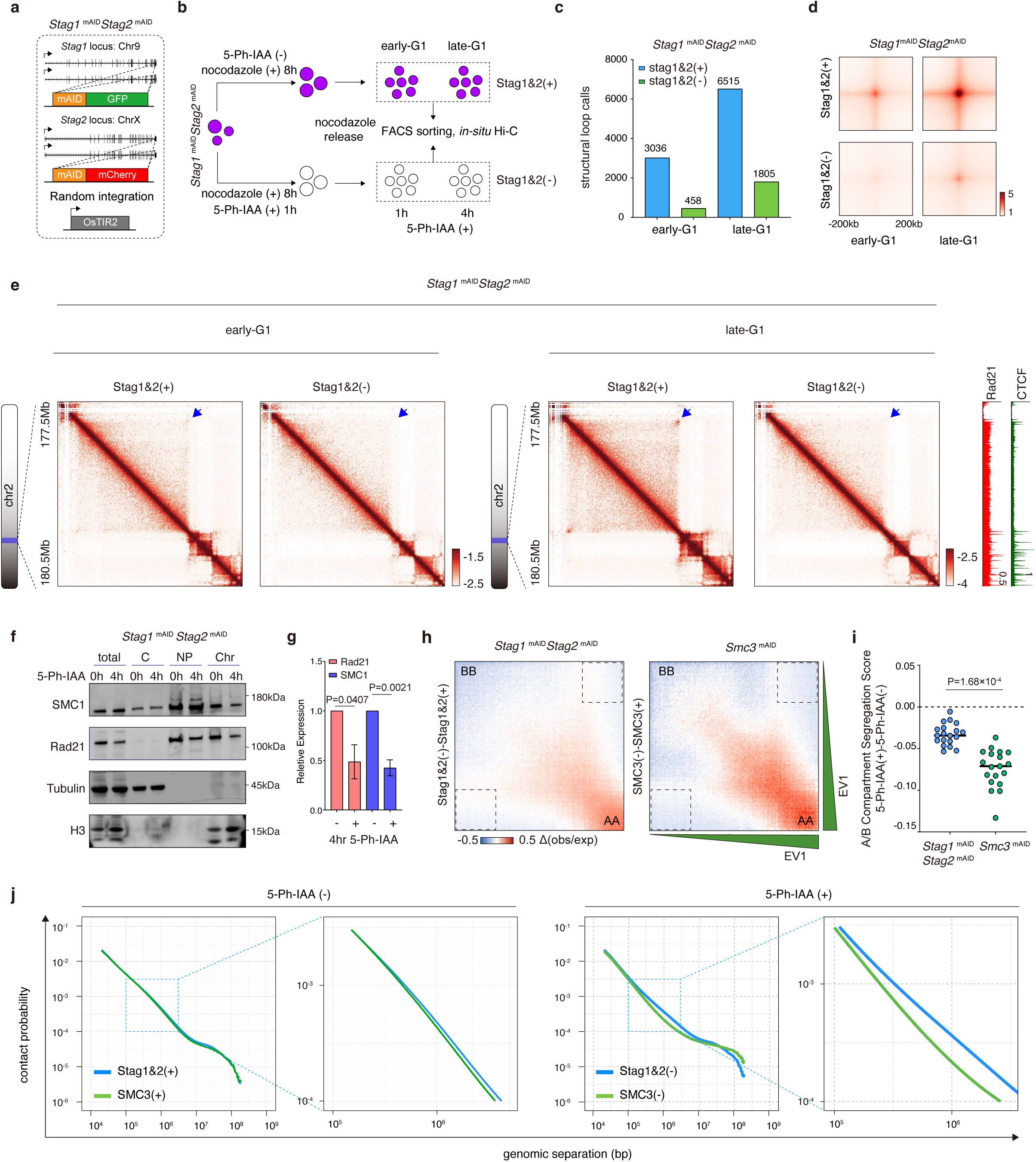
Impacts of Stag1 and Stag2 co-depletion on post-mitotic chromatin structural re-configuration. **a**, Schematic of the *Stag1*^mAID^ *Stag2*^mAID^ line: the Stag1 locus carries an mAID-GFP tag and the Stag2 locus carries an mAID-mCherry tag, with an OsTIR2 transgene ectopically expressed. **b**, Workflow for mitotic arrest/release and 5-Ph-IAA treatment to obtain early-G1 and late-G1 populations with or without Stag1 and Stag2. **c**, Bar graph showing the number of structural loop calls in early- and late-G1 cells with or without Stag1 and Stag2. **d**, APA plots showing the massive loss of Structural loops after mitosis upon Stag1 and Stag2 co-depletion. **e**, Representative Hi-C contact maps showing structural loop loss (blue arrows) after mitosis upon Stag1 and Stag2 co-depletion. Late-G1 CTCF and Rad21 ChIP-seq tracks are shown on the right. **f**, Representative immunoblots of chromatin fractionation experiment showing that chromatin bound cohesin subunits are reduced but not completely abolished upon Stag1 and Stag2 co-depletion (Total, total protein; C, cytoplasm; NP, nucleoplasm; Chr, chromatin). Three independent experiments were performed. **g**, Quantification of (**f**) showing that ∼50% of Rad21 and SMC1 is retained upon Stag1 and Stag2 co-depletion. Error bar denotes mean ± s.d. n=3 independent experiments. *P* values were calculated using two-tailed Student’s t-test. **h**, Subtracted Saddle plots showing that Stag1 and Stag2 co-depletion induced less pronounced changes in chromatin compartmentalization compared to SMC3 depletion (dotted boxes). **i**, Dot plots showing that Stag1 and Stag2 co-depletion strengthened A/B compartment segregation. Yet, the effect size was smaller compared to that induced by SMC3 loss. Each dot represents a single chromosome. *P* values were calculated using two-sided, paired Wilcoxon rank-sum tests. **j**, *P(s)* curves showing higher short-range (loop and TAD scale) contact frequency in Stag1 and Stag2 co-depleted cells compared to SMC3-deficient cells, indicating residual loop extrusion activity.

HICCUPS analysis revealed a massive loss of structural loops after mitosis, in the absence of both Stag1 and Stag2 (Fig. 5c). In line with this, visual examination of Hi-C contact matrices at individual genomic loci and genome-wide APA plots both unveiled a drastic elimination of structural loop signals in both early- and late-G1 phase (Fig. 5d, e). Taken together, our data demonstrate that at least one Stag protein is required for post-mitotic loop reformation. Although the loss of either Stag1 or Stag2 alone induces only marginal defects, their combined ablation synergistically prevents the re-establishment of the loop architecture, demonstrating their essential and redundant roles in this process.

### Co-depletion of Stag1 and Stag2 impairs chromatin association of cohesin without affecting its loop extrusion processivity

To investigate the mechanism through which co-depletion of Stag1 and Stag2 abrogates structural loops, we sought to determine if this failure resulted from impaired cohesin chromatin binding or reduced loop extrusion processivity. Chromatin fractionation revealed a significant reduction in chromatin-bound cohesin core subunits (SMC1 and Rad21) after four hours of 5-Ph-IAA treatment, confirming compromised cohesin loading (Fig. 5f, g). Of note, approximately half of the cohesin remained chromatin-associated, suggesting that a population of cohesin complexes can load in a Stag1- and Stag2-independent manner (Fig. 5f, g).

Next, to assess whether the remaining chromatin-bound cohesin complexes retained loop extrusion capacity, we established a comparative framework using targeted depletion of two key chromatin structural modulators: the cohesin core subunit SMC3 and the architectural protein CTCF. Depletion of SMC3 served as a benchmark for complete extrusion failure and caused a profound loss of TAD/loop scale (100kb-1Mb) contacts on the *P_S_* curve, a complete disruption of structural loops, and sharply enhanced A/B compartmental segregation (Fig. 5h; Extended Data Fig. 9a–d). Conversely, CTCF loss, which eliminates structural loops but retains full extrusion capacity, failed to induce any changes in the *P_S_* curve or compartmentalization (Extended Data Fig. 9e-h). These results establish that changes in the *P_S_* curve and compartment strength are sensitive and specific readouts of cohesin-mediated loop extrusion activity.

Co-depletion of Stag1 and Stag2 produced changes in the *P_S_* curve and compartmentalization that are qualitatively similar to those of SMC3 deficiency, confirming impaired loop extrusion (Fig. 5h, i; Extended Data Fig. 9i). However, the magnitude of these effects was substantially smaller than that in SMC3-deficient cells (Fig. 5h–j), indicating that extrusion activity was attenuated but not completely abolished. Collectively, these results position the Stag1 and Stag2 co-depletion phenotype as intermediate between complete extrusion loss (SMC3 loss) and full extrusion activity with loop dissolution (CTCF loss). This partial impairment strongly suggests that the residual population of chromatin-bound cohesin retains DNA loop extrusion capability even in the absence of both Stag subunits.

We next asked whether Stag1 and Stag2 loss also compromised extrusion processivity—that is, the distance that cohesin could potentially travel along DNA—of the remaining complexes. We found that in contrast to Nipbl-deficient cells, which exhibit a preferential loss of large loops due to impaired cohesin extrusion processivity ^37,38^, Stag1 and Stag2 co-depletion uniformly reduced structural loops across all size ranges (Extended Data Fig. 10a-c). This lack of size-dependent attenuation indicates that the intrinsic extrusion processivity of chromatin-bound cohesin remains intact without Stag1 and Stag2. The global reduction in loop strength likely results from decreased cohesin chromatin occupancy.

### Stag1 and Stag2 distinctly influence post-mitotic gene reactivation

The distinct impacts of Stag1 and Stag2 on genome reorganization prompted us to investigate their respective roles in gene reactivation following mitosis. To approach this, we utilized TT-seq to measure nascent transcription during early- and late-G1 phases in cells acutely depleted of either Stag1 or Stag2^39^. Notably, Stag1 depletion failed to induce any significant transcriptional changes, which aligns with its negligible effect on post-mitotic genome refolding (Fig. 6a, b; Extended Data Fig. 11a). In contrast, Stag2 deletion triggered subtle but reproducible transcriptional changes after mitosis, yielding a total of 35 differentially expressed genes (DEGs, Padj < 0.05, fold change > 1.5), with 17 and 30 DEGs identified in early- and late-G1 phases, respectively (Fig. 6c, d; Extended Data Fig. 11b). Among these, 17 and 26 DEGs were down-regulated in early- and late-G1 phases, respectively, suggesting that Stag2 primarily participates in transcription activation. Of note, although the late-G1 phase displayed more DEGs than early-G1 (30 and 17 genes respectively), the magnitude of transcriptional changes (log_2_FC) was comparable between these two stages (Fig. 6e), suggesting that Stag2 contributes to rapid transcription initiation after mitosis. This finding is consistent with our prior finding that Stag2 is essential for the swift formation of small structural loops, which likely facilitates the rapid establishment of promoter-enhancer (P-E) contacts after mitosis.

**Figure 6:**
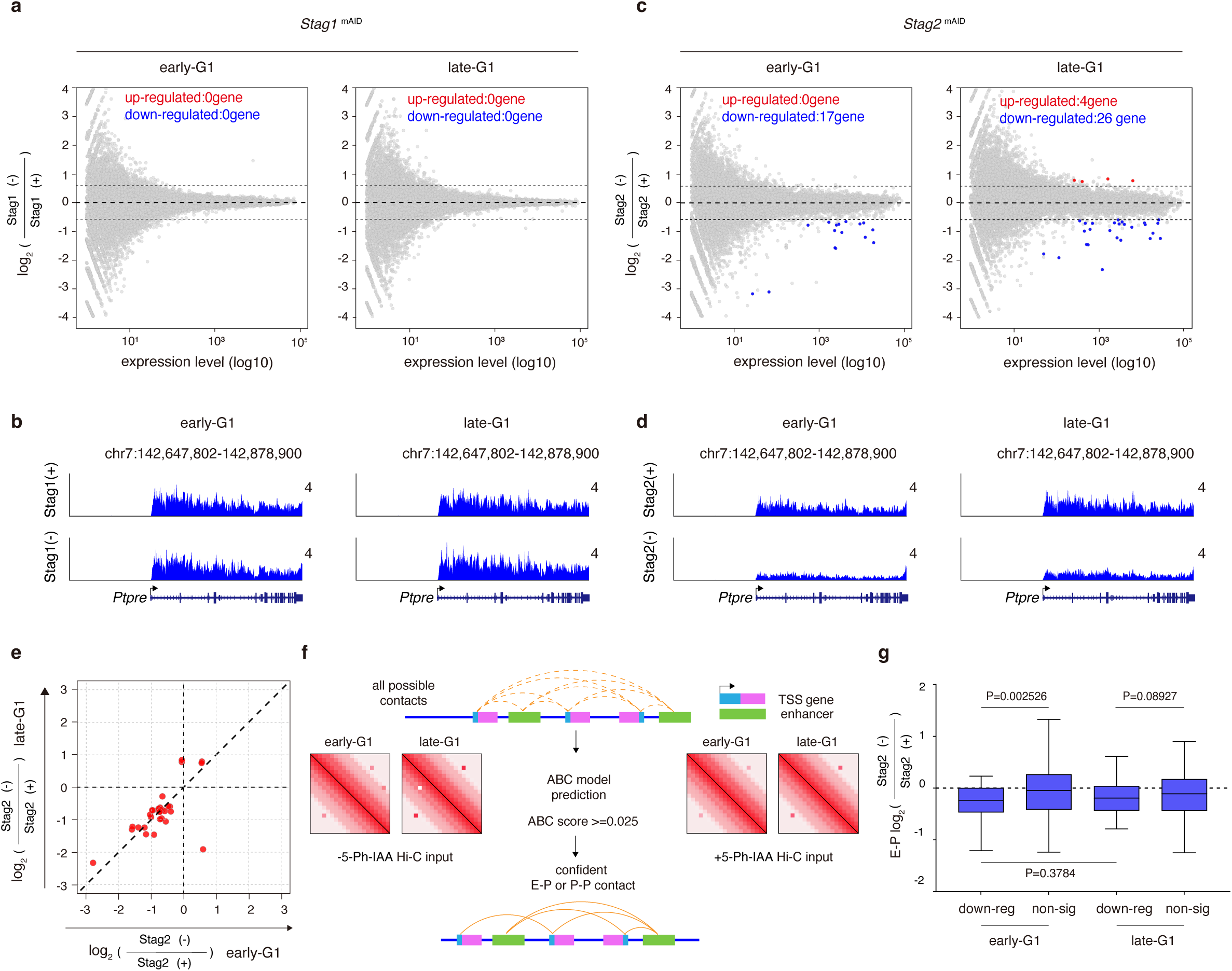
Stag1 and Stag2 diversely affect post-mitotic gene reactivation. **a,** Scatter plots showing nascent transcription changes upon Stag1 depletion in early- and late-G1 phase. **b**, Genome browser tracks at the *Ptpre* locus showing TT-seq signals with or without Stag1 depletion in early- and late-G1 phase. **c**, Scatter plots showing nascent transcription changes upon Stag2 depletion in early- and late-G1 phase. **d**, Genome browser tracks at the *Ptpre* locus showing TT-seq signals with or without Stag2 depletion in early- and late-G1 phase. Note that *Ptpre* is selectively affected by Stag2 loss. **e**, Scatter plot showing comparable log_2_FC of down-regulated genes between early- and late-G1 phase upon Stag2 loss. **f**, Schematic of the ABC model workflow predicting P-E contacts using Hi-C input with or without Stag2. **g**, Boxplots showing log_2_FC in P-E contact strength for down-regulated and non-significant genes upon Stag2 depletion in early- and late-G1 phases. *P* values for comparisons between down-regulated and non-significant genes were calculated using the two-sided Wilcoxon rank-sum test. Paired Wilcoxon test was used to compare the changes in the same set of down-regulated genes between early- and late-G1 phases.

To determine if the reduced transcription upon Stag2 loss stemmed from dysregulated P-E interaction, we applied the activity-by-contact (ABC) model to identify high-confidence P-E contacts (Fig. 6f). Using an ABC cutoff of 0.025, we identified 86 P-E contacts, associated with 27 down-regulated genes. Importantly, these contacts were dramatically attenuated upon Stag2 depletion with comparable magnitudes between early- and late-G1 phase cells (Fig. 6g). In contrast, P-E contacts associated with non-differentially expressed genes were not as severely affected (Fig. 6g). These results indicate that Stag2 enables rapid post-mitotic reactivation of specific genes by mediating functional P-E interactions.

### Co-depletion of Stag1 and Stag2 results in a more robust change in post-mitotic transcription

Given the profound synergistic impairment of post-mitotic genome refolding upon Stag1 and Stag2 co-depletion, we asked whether their combined loss also synergistically disrupts transcription reactivation. Indeed, deleting both Stag proteins elicited a markedly more pronounced effect on primary transcript levels compared to individual depletion of either Stag1 or Stag2, yielding 90 and 760 DEGs in early- and late-G1 phase, respectively (Fig. 7a). Among the total 792 DEGs, 547 genes were significantly downregulated, confirming a preferential role of Stag proteins in activating transcription. In contrast to the rapid transcriptional dysregulation caused by Stag2 depletion alone, simultaneous loss of Stag1 and Stag2 resulted in a progressively more severe disruption that became significantly more pronounced by late-G1 phase (Fig. 7b). To further investigate this temporal response pattern of dysregulation, we stratified the downregulated genes into three groups, based on the timing of their response: early-, intermediate-, and late-affected genes (Fig. 7b, c; Extended Data Fig. 11c). Notably, early-affected genes were more frequently associated with enhancers (∼68%) compared to intermediate-(∼52.6%), late-(∼52.7%), or non-affected (∼18%) genes (Fig. 7d), implying a stronger dependence on distal regulatory elements. Furthermore, the early-affected genes exhibited weaker H3K27ac signals at promoters and lower expression levels compared to intermediate- and late-affected groups (Fig. 7e-g; Extended Data Fig. 11d). These features are consistent with a recent report that weakly active promoters (often linked to tissue-specific genes) lack intrinsic enhancer-like sequences and rely more heavily on P-E contacts for their expression, whereas highly active promoters (often linked to housekeeping genes) usually harbor built-in enhancer elements and are less reliant on distal enhancers^40^. Interestingly, enhancers associated with each group of genes displayed comparable levels of H3K27ac marks (Fig. 7e, f).

**Figure 7:**
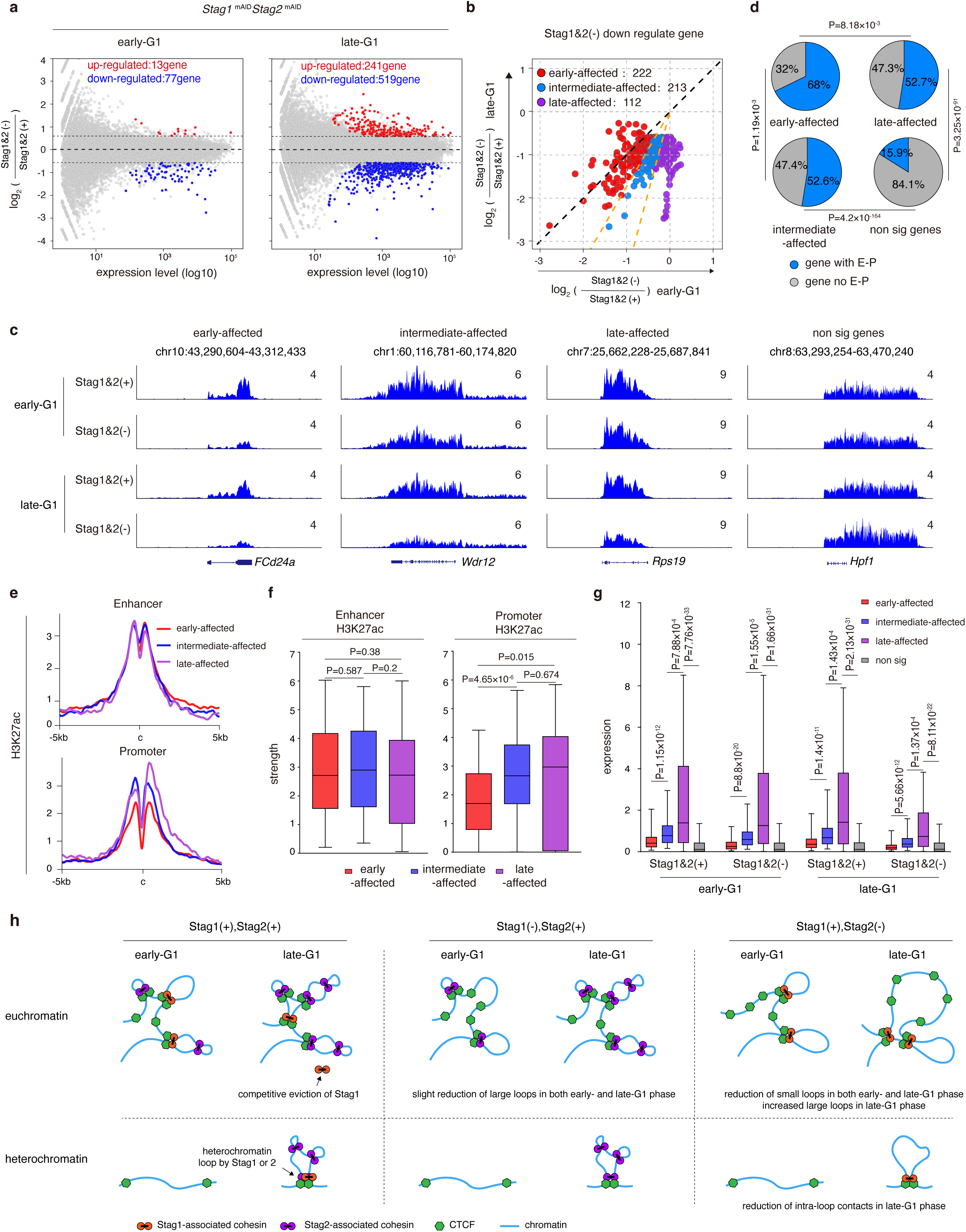
Co-depletion of Stag1 and Stag2 induces a robust post-mitotic transcriptional change. **a**, Scatter plots showing the changes in nascent transcription upon Stag1 and Stag2 co-depletion in early- and late-G1 phase. **b**, Scatter plot comparing log_2_FC values of down-regulated genes upon Stag1 and Stag2 co-depletion, between early- and late-G1 phase. **c**, Genome browser tracks showing TT-seq signal of *Fcd24a* (early-affected), *Wdr12* (intermediate-affected), *Rps19* (late-affected) and *Hpf1* (non-significant) loci upon Stag1 and Stag2 co-depletion in early- and late-G1 phase. **d**, Pie charts showing the percentage of genes with predicted P-E contacts (ABC score ≥ 0.025) across indicated groups. *P* values are determined using two-sided Fisher’s exact tests. **e**, Metaplots of H3K27ac ChIP-seq signal at enhancers linked to the indicated gene sets (top) and at promoters these genes (bottom). **f**, Boxplots showing the quantification of (**e**). *P* values are calculated using two-sided Wilcoxon rank-sum test. **g**, Boxplots showing TT-seq signals in the gene body for early-affected, intermediate-affected, late-affected and non-significant gene groups. Data are shown for early- and late-G1 cells, with or without Stag1 and Stag2. *P* values are calculated using a two-sided Wilcoxon rank-sum test. **h**, Schematic summarizing the distinct and compensatory roles of Stag1 and Stag2 during post-mitotic genome folding. Stag1 and Stag2 are dynamically loaded into euchromatin after mitosis, with Stag2’s high nuclear concentration in late-G1 competitively restricting Stag1 chromatin association (top left panel). Stag1 depletion mildly reduces large loops (top middle panel), whereas Stag2 depletion reduces small loops throughout G1 and promotes Stag1-mediated large loop formation in late-G1 (top right panel). Stag2-mediated dynamic loop extrusion in heterochromatin is delayed until late-G1 (bottom panels).

Taken together, our data indicate that the distinct timing of gene dysregulation following Stag loss may stem from their differential reliance on distal enhancers. Specifically, genes with weak promoters heavily rely on the rapid re-establishment of P-E contacts after mitosis to ensure efficient transcription, a process facilitated by Stag1- or Stag2-associated cohesin.

## Discussion

### Distinct impacts of Stag1 and Stag2 loss on genome refolding

Our study systematically dissected the individual and combinatorial contributions of Stag1 and Stag2 to post-mitotic genome refolding and transcription reactivation. Based on Stag1’s rapid post-mitotic chromatin recruitment, we hypothesized that Stag1-associated cohesin would mediate initial small-loop formation and predicted that structural loop re-establishment would be delayed in its absence^25^. However, to our surprise, genome re-configuration was largely unaffected by Stag1 loss, with only minor defects observed in large loop reformation (Fig. 7h top left and top middle panels). This unexpected insensitivity of structural loops to Stag1 loss highlights Stag2’s full compensatory capacity and is consistent with its status as the dominant Stag paralog^36,41^. Furthermore, this finding indicates that despite Stag2’s progressive pattern of post-mitotic recruitment, its initial loading in early-G1 is sufficient to establish most structural loops (Fig. 7h top left and top middle panels). Stag1 depletion also induced no measurable changes in post-mitotic transcriptional reactivation, in line with its limited impacts on genome refolding. This result mildly contrasts with prior reports of transcriptional alterations upon Stag1 loss, a discrepancy that may reflect cell type-specific differences or technical variations^34,36^. Nevertheless, Stag1’s overall impact on genome organization consistently appears smaller than that of Stag2 across various studies, aligning with the established clinical observation that Stag2 mutations significantly outnumber Stag1 in human diseases, including many types of cancer^41-44^. Our findings reveal a crucial dual role of Stag2 in post-mitotic loop formation. Stag2 loss induced an immediate and persistent weakening of small structural loops (<100 kb) throughout the entire G1 phase (Fig. 7h top right panel). Conversely, large loops were strengthened upon Stag2 loss, aligning with previous reports (Fig. 7h top right panel)^20^. This effect is specifically observed in late-G1 phase, consistent with Stag2’s progressive nuclear import. As Stag2’s nuclear concentration gradually increases in late-G1, it reaches a level high enough to compete with Stag1 for cohesin subunits (Fig. 7h top left panel). Notably, the absence of large loop suppression by Stag2 in early-G1 suggests that at this stage, the nuclear pool of core cohesin subunits is sufficient to accommodate both Stag proteins (Fig. 7h top left panel).

### Factors affecting Stag2’s loop extrusion capacity

In late-G1, Stag2 competes with Stag1 for cohesin core subunits but fails to support large-loop formation. This functional deficiency may arise from two biophysical constraints: First, distance-dependent limitations, evidenced by progressively stronger suppression at larger sizes (>300 kb); Second, intermediate CTCF barriers that may actively terminate Stag2-mediated extrusion. Our analysis revealed that the presence of such obstacles significantly attenuates Stag2’s loop-forming capacity. Specifically, each additional barrier markedly increases dissociation events. Beyond a critical threshold (typically two or more intermediate CTCF barriers), the influence of genomic distance becomes negligible. This finding underscores that physical barriers supersede travel distance as the primary determinants restricting the sizes of Stag2-mediated loops, effectively confining Stag2 to smaller, less obstructed genomic regions.

### A dynamic and chromatin context-dependent regulation of Stag2 in post-mitotic genome refolding

Our data revealed a dynamic and chromatin-context dependent loading of Stag2 after mitosis, preferentially associating with euchromatin in early-G1, followed by a delayed onset of heterochromatin recruitment in late-G1 (Fig. 7h bottom panels). In heterochromatin, we observed a gradual intensification of intra-loop contacts in control cells that were notably attenuated in Stag2-deficient cells. These intra-loop contacts likely represent extrusion intermediates not yet anchored by CTCF. Conversely, in euchromatin, Stag2 depletion promptly induced a loss of intra-loop contacts in early-G1, suggesting a preferential recruitment to these regions after mitosis (Fig. 7h top left and top right panels). This swift association likely facilitates efficient post-mitotic transcription reactivation by enabling rapid re-linking between enhancers and promoters through dynamic extrusion intermediates. The precise mechanism underlying the preferential loading of Stag2 to euchromatin over heterochromatin remains to be elucidated. We speculate that Stag2-associated cohesin is actively recruited to open chromatin via engagement with active cis-regulatory elements, whereas its incorporation into heterochromatin may occur through a more passive or delayed mechanism.

### Stag proteins regulate cohesin-chromatin association

Cohesin loading is widely attributed to Nipbl, which exhibits high DNA-binding affinity^8,45-48^. It is therefore surprising that depletion of Stag proteins extensively reduces cohesin loading and loop formation. This effect is unlikely to stem from impaired holoenzyme assembly, as the cohesin trimer forms stable complexes even in the absence of DNA, ATP, or accessory subunits^49^. Instead, our findings align with recent biochemical evidence that Stag1 significantly enhances cohesin’s DNA binding affinity^49^. The marked reduction in chromatin-associated cohesin upon Stag depletion can be explained by the loss of this cooperative DNA-binding activity^49^. The residual cohesin retention in Stag1- and Stag2-deficient cells may reflect compensatory tethering by Nipbl, which retains autonomous DNA binding capacity. Nevertheless, our results underscore that Stag proteins play an important role in cohesin-chromatin interactions and loop extrusion.

In summary, our study in synchronized post-mitotic cells reveals a functional division of labor between Stag1 and Stag2 during genome refolding. Stag2 acts as the primary regulator, playing a critical role in the formation of small loops in early-G1, a process that is important for efficient transcriptional reactivation after mitosis. It also competitively restricts large loop formation in late-G1. In contrast, Stag1 provides supportive and compensatory functions, contributing modestly to larger loop formation and serving as a failsafe mechanism that maintains genome architectural integrity in the absence of Stag2. This cooperation ensures robust transcriptional regulation and could prevent catastrophic structural failure after mitosis, underscoring the essential yet distinct contributions of both Stag paralogs to cohesin-mediated genome organization.

## Acknowledgements

We thank Nicolas G. Aboreden and members of the Zhang lab for helpful discussions. We acknowledge the technical support from Zhiqing Huang, Mengyuan Li, and Dechun Cheng at the Flow Cytometry Core of Shenzhen Bay Laboratory. This work was supported by the National Natural Science Foundation of China (grant no. 82471197 to H. Zhang), the Shenzhen Medical Research Fund (grant no. D2401002 to H. Zhang), and the Major Program of Shenzhen Bay Laboratory (grant no. S201101004 to H. Zhang).

## Author contributions

H.Z. and M.W. conceived the study and designed experiments. Manzhu Wang performed experiments with help from F.S., B.Z., D.J., L.S. Data analysis was performed by M.W. with help from S.X., L.S., F.L., Y.W., and B.L. H.Z. and M.W. wrote the manuscript with inputs from all authors.

## Methods

### Cell culture

The parental G1E-ER4 cell line was kindly provided by the laboratory of Dr. Mitchell Weiss. G1E-ER4 cells were cultured in suspension in IMEM medium supplemented with 13% fetal bovine serum (FBS, v/v), 0.2mM penicillin-streptomycin (Gibco), 179ng/ml murine stem cell factor, 3.47ng/ml Epogen, and 0.001% 1-thioglycerol (v/v). The cell density was maintained below 1×10^6^ cells/ml. Sublines of G1E-ER4 cells expressing mAID-GFP-tagged Stag1, mAID-mCherry-tagged Stag2, or both tags were maintained under the same culture conditions as the parental line. All cell lines were cultured in a humidified incubator at 37°C with 5% CO_2_. All cell lines were routinely tested and confirmed to be mycoplasma-negative.

### CRISPR/Cas9 mediated genome editing

To generate *Stag1*^mAID^ and *Stag2*^mAID^ knock-in cell lines, ∼800 bp homology arms flanking the stop codons of the *Stag1* and *Stag2* genes were PCR-amplified from the genomic DNA of G1E-ER4 cells and cloned into the pcDNA3.1 vector. The coding sequences of EGFP-mAID (for Stag1) or mCherry-mAID (for Stag2) were inserted immediately upstream of the stop codon to construct homology-directed repair (HDR) donor plasmids. Single guide RNAs (sgRNAs) targeting the 3′ ends of the *Stag1* and *Stag2* genes were designed using the Benchling online tool and cloned into the pX330 vector. G1E-ER4 cells were electroporated with 18µg of the HDR donor plasmid and 18µg of sgRNA-containing pX330 plasmid using an Amaxa 2b Nucleofector (Lonza) with program G-016. 24 to 48 hours post-electroporation, single cells with positive EGFP or mCherry signals were sorted into 96-well plates using FACSAria III (BD Biosciences). Clones were expanded and genotyped by PCR using primers flanking the integration site. Positive clones were further verified by Sanger sequencing.

To generate the double knock-in *Stag1*^mAID^ *Stag2*^mAID^ cell line, *Stag2*^mAID^ cells were electroporated with donor and sgRNA containing pX330 plasmids corresponding to *Stag1* using Amaxa 2b Nucleofector (Lonza) with program G-016. Genotyping and validation were performed as described above.

To enable rapid degradation of Stag1 or/and Stag2, OsTIR2 was introduced into *Stag1*^mAID^, *Stag2*^mAID^, and *Stag1*^mAID^ *Stag2*^mAID^ cells as previously described^24,33,50,51^. Briefly, a retroviral vector encoding an OsTIR2-TagBFP fusion protein was transfected into co-HEK293T cells together with pCL-Eco packaging plasmid. Virus-containing supernatant was collected 48 and 72 hours later. Target cells were infected using a spin infection protocol, which included polybrene and HEPES. 48 hours after infection, cells expressing the TagBFP protein were isolated by sorting using a BD FACSAria III.

### Cell growth curve

To examine the impact of Stag1 and/or Stag2 depletion on cell growth, 50,000 *Stag1*^mAID^, *Stag2*^mAID^ or *Stag1*^mAID^*Stag2*^mAID^ cells were seeded into untreated non-adhesive 6-well plates with 2ml of fresh culture medium per well. Cells were then treated with or without 1μM 5-Ph-IAA. Cell numbers for each condition were counted every 24 hours over a total period of 96 hours. Three independent biological replicates were performed for each condition.

### Western Blotting

Western blotting was performed as previously described^51^. 3×10^6^ cells were lysed with RIPA buffer (Beyotime, p0013b) supplemented with 1:100 protease inhibitor (Sigma-Aldrich, P8340) on ice for 20 minutes. Cell lysates were sonicated using a QSonica Q800R3 sonicator (40% amplitude, 5 seconds ON and 5 seconds OFF) for 5 minutes. Blots were probed with anti-GFP antibody (ABclonal, AE012, 1:5000) and anti-mCherry antibody (Abcam, ab167453, 1:5000), followed by imaging using a BioRad ChemiDoc MP Imaging System.

For chromatin fractionation, approximately 1×10^7^ cells were harvested. 20% of cells were saved as the whole cell lysate(total). The remaining cells were washed once with PBS, pelleted (500g, 3 min, 4°C), and resuspended in Buffer A (10mM HEPES pH 7.9, 10mM KCl, 1.5mM MgCl_2_, 0.34M sucrose, 10% glycerol, 0.4% Triton X-100, supplemented with 0.5 mM PMSF, 1μg/ml aprotinin, 1μg/ml leupeptin, and 1μg/ml pepstatin). After incubation on ice for 10min, the mixture was rotated at 4°C for 1 hour, lysates were centrifuged (1300g, 4 min, 4°C). The supernatant was collected as the cytoplasmic fraction (cytoplasm, C). The pellet (nuclei) was washed three times with detergent free Buffer A and then resuspended in Buffer B (3mM EDTA, 0.2mM EGTA, 300mM NaCl, 1mM DTT, 0.3% NP-40, supplemented with 1mM PMSF, 1μg/ml aprotinin, 1μg/ml leupeptin, and 1μg/ml pepstatin) and incubated on ice for 5 min. Samples were centrifuged (1700g, 4 min, 4°C) to separate the supernatant (nucleoplasm, NP) and pellet (chromatin, Chr). Total and P3 fractions were further lysed in nuclei lysis buffer (50mM Tris-HCl pH 8.0, 10mM EGTA, 1% SDS) and sonicated (Qsonica Q800R3; 40% amplitude, 10s ON / 5s OFF, 6min total). Equal protein amounts from Total, S2, S3, and P3 fractions were mixed with 5×SDS loading buffer, boiled at 100°C for 10min, centrifuged (13,000g, 1min), and subjected to SDS–PAGE and immunoblotting.

### Nocodazole mediated prometaphase arrest/release and protein degradation

To examine post-mitotic chromatin re-configuration in the absence of Stag1 and/or Stag2, *Stag1*^mAID^, *Stag2*^mAID^ or *Stag1*^mAID^*Stag2*^mAID^ cells were arrested at prometaphase using nocodazole (200ng/ml) for 8 hours. Toward the end of nocodazole arrest, 5-Ph-IAA (1μM) was applied to these cells for 1 hour. After nocodazole treatment, cells were released into warm fresh medium containing 5-Ph-IAA but not nocodazole for 1 hour (early-G1) and 4 hours (late-G1), respectively. Cells without auxin treatment were processed in parallel as controls.

### In-situ Hi-C

*In-situ* Hi-C was performed as previously described^51^. Briefly, formaldehyde-fixed cells (10^5^) were lysed on ice. Nuclei were isolated, washed, and chromatin was opened with SDS at 65°C for 10 min. SDS activity was quenched with Triton X-100, followed by incubation at 37°C for 1 hour. Chromatin was digested overnight with DpnII (25 U) at 37°C, with an additional 25 U added for 2 hours the next day. DpnII was inactivated at 65°C for 30min. DNA ends were filled-in using Klenow fragment, with dGTP, dCTP, dTTP and biotin-14-dATP at 37°C. Proximity ligation was performed with T4 DNA ligase (2000 U) at 16°C (4h) and then 25°C (2h). Crosslinks were reversed overnight with proteinase K and SDS at 65°C, followed by RNase digestion. DNA was extracted and sonicated (15s ON / 15s OFF, 12.5 min total). Biotinylated fragments were captured using pre-washed Dynabeads MyOne Streptavidin beads. Beads were washed sequentially with 1×B&W buffer and T4 ligase buffer. End repair, adaptor ligation and bead washing were performed using Vazyme general MGI library construction kit (N607). Bound DNA was eluted in 0.1% SDS at 98°C and purified using 1.8× VAHTS DNA Clean Beads (N411-02). DNA was PCR amplified (9 cycles), purified 0.9× VAHTS DNA Clean Beads, and quantified by Qubit. Final libraries were sequenced on the MGI DNBSEQ-T7 platform.

### Chromatin immunoprecipitation and sequencing (ChIP-seq)

ChIP-seq experiments were performed as previously described^51^. Briefly, for each IP, 1×10^7^ cells were lysed in cold cell lysis buffer (10mM Tris–HCl pH 8.0, 10mM NaCl, 0.2% NP-40) with protease inhibitor (Sigma-Aldrich, P8340) and PMSF (Sangon Biotech, A100754) on ice for 20 minutes. Nuclei were then lysed in nuclear lysis buffer (50mM Tris–HCl pH 8.0, 10mM EDTA, 1% SDS) with PI/PMSF on ice for 20 minutes. Chromatin was sonicated using a QSonica Q800R3 (80% amplitude, 20s ON/40s OFF, 16 min total). The sonicated chromatin was diluted with 4 volumes of IP dilution buffer and pre-cleared for 4-6 hours at 4°C with 25μl protein A/G agarose beads (Santa Cruz, sc-2003). The pre-cleared chromatin was then incubated overnight at 4°C with beads bound to antibodies against Stag1 (Abcam, ab4455) or Stag2 (CST, 5882). Following the overnight incubation, the beads were washed sequentially with IP wash buffer 1 (20 mM Tris pH 8, 2mM EDTA, 50mM NaCl, 1%Triton X-100, 0.1% SDS), high-salt buffer (20 mM Tris pH 8, 2mM EDTA, 500mM NaCl, 1% Triton X-100, 0.01% SDS), IP wash buffer 2 (10mM Tris pH 8, 1mM EDTA, 0.25M LiCl, 1% NP-40/Igepal, 1% sodium deoxycholate), and TE buffer. The bound DNA was eluted in 200μl elution buffer (100mM NaHCO₃, 1% SDS) for 5 minutes at 65°C. Crosslinking was reversed by incubating the samples overnight at 65°C. The samples were then treated with 2μl RNase A (10mg/ml) at 37°C for 30 minutes, followed by 3μl proteinase K (20mg/ml) at 65°C for 2 hours. Finally, the DNA was purified using an Omega PCR clean up kit, eluted in 60μl of water, and prepared for sequencing using the VAHTS® Universal DNA Library Prep Kit for Illumina V3 (Vazyme ND610). The libraries were then sequenced on the MGI DNBSEQ-T7 platform.

### Transient transcriptome sequencing (TT-seq)

Cells were synchronized and released based on the aforementioned nocodazole arrest/release protocol. To metabolically label newly synthesized RNA, 4-thiouridine (4sU; Sigma Aldrich, T4509) was added to the cell culture medium to a final concentration of 500µM. Cells were incubated at 37°C for 10 minutes before harvesting. Approximately 1×10^7^ cells were collected at 1h and 4h after mitosis. Total RNA was extracted using TRIzol reagent (Thermo Fisher Scientific, 15596026CN), followed by chloroform extraction and isopropanol precipitation. The resulting RNA pellet was washed twice with 75% ethanol, air-dried, and resuspended in RNase-free water. The RNA was then fragmented using NaOH, immediately followed by neutralization with Tris-HCl (pH 6.8), DTT (1 mM; Sigma Aldrich, 43815), and sodium acetate (pH 5.5). The fragmented RNA was purified by isopropanol precipitation. Fragmented RNA was biotinylated by incubating it with EZ-Link HPDP-Biotin (Thermo Fisher Scientific, 21341) with gentle rotation. After chloroform extraction and isopropanol precipitation, the biotinylated RNA was captured using Dynabeads MyOne Streptavidin C1 (Thermo Fisher Scientific). To remove unbound RNA, the beads were washed sequentially with a 1×B&W buffer (200µl) and high-salt buffer (200µl). The captured RNA was eluted with 100 mM DTT, purified, and then resuspended in RNase-free water. Strand-specific RNA-seq libraries were prepared from 100 ng of the purified RNA using the VAHTS® Universal V8 RNA-seq Library Prep Kit for MGI (Vazyme, NRM605), following the manufacturer’s protocol. The prepared libraries were then sequenced on the MGI DNBSEQ-T7 platform.

### Quantification and data analysis

#### ChIP-seq

Paired-end reads were first aligned to the human reference genome (hg19) using Bowtie2 (v2.3.5.1) with default parameters and soft clipping enabled (“--local”). Reads that did not map to the human genome were subsequently aligned to the mouse reference genome (mm9). Low-quality alignments (MAPQ < 30), PCR duplicates, and reads mapped to mitochondrial DNA, unplaced contigs, or ENCODE blacklisted regions were removed using SAMtools (v1.9) and BEDTools (v2.27.1). For data normalization, reads were processed using the bamCoverage function from deepTools (v3.1.3) with the normalization method set to counts per million mapped reads (CPM).

#### Peak calling

Peak calling was conducted separately for each biological replicate using MACS2 (v2.2.7.1) with default parameters and a q-value cutoff of 0.01 for narrow peaks. The corresponding input samples were used as background controls for MACS2 peak calling.

#### Hi-C data preprocessing

Hi-C data processing was performed as previously described^24,33,50,51^. Briefly, Hi-C data from each biological replicate and replicate-merged samples were processed using the HiC-Pro (v3.0.0) pipeline. Sequencing reads were first mapped to the mouse reference genome (mm9) using Bowtie2 (v2.3.5.1) with optimized global and local alignment parameters (For global alignment, the parameters were --very-sensitive -L 30 --score-min L, -0.6,-0.2 --end-to-end --reorder, and for local alignment, the parameters were --very-sensitive -L 20 --score-min L, -0.6,-0.2 --end-to-end --reorder). The resulting alignments were then filtered to identify valid interaction pairs by removing invalid ligation products, self-ligated fragments, and PCR duplicates. The filtered pairs were subsequently converted into .hic and .cool file formats for visualization with Juicebox and for quantitative analysis with HiCExplorer (v3.7), respectively. All contact maps, saddle plots, and aggregate peak analysis (APA) plots in the main figures were generated from merged datasets of all biological replicates.

#### Eigenvector decomposition and saddle plots

Compartmentalization analysis was performed as previously described^24,50,51^. Briefly, eigenvector decomposition was used to identify A/B compartments. The first eigenvector (EV1) was derived from a Pearson’s correlation matrix from a 25kb-binned, Knight-Ruiz (KR) balanced cis-interaction matrices. The orientation of the EV1 was determined based on gene density. Following this, saddle plots were generated to assess chromatin compartmentalization strength. The 25kb-binned, KR-balanced observed/expected *cis*-contact matrices were extracted using the DUMP function of Juicer Tools. These matrices were then reordered based on the ascending order of EV1 values from asynchronous sample. This reordering places A-A and B-B interactions at the lower-right and upper-left corner respectively. The reordered contact maps were then subdivided into 200×200 blocks, and the interaction frequencies were averaged to create genome-wide saddle plots. Compartment strength for each chromosome was quantified as the ratio of within-compartment interactions to inter-compartment interactions:

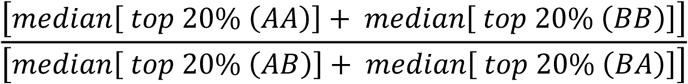

All saddle plots in the main figures were generated from merged biological replicates.

#### Contact probability decay curves (*P(s)* curves)

For each sample, the contact probability decay curve *P(s)* was calculated using the “expected-cis” function of Cooltools (v0.4.0).

#### Loop calling

Loop calling was performed as previously described^50,51^. Briefly, preliminary loops were identified from KR-balanced contact maps using the HICCUPS function in juicer_tools (v1.22.01) at two resolutions: 10kb and 25kb. Specific parameters were used for each resolution: for 10 kb, the parameters were p = 4 bins, i= 16 bins, and an FDR of 0.2, while for 25kb, the parameters were p = 5 bins, i = 15 bins, and an FDR of 0.1. Because redundant loops can be called across different samples, a greedy clustering algorithm was used to merge them. Loops were first ranked in ascending order based on their minimum q-value across all samples. Starting with the top-ranked loop, a search was performed for nearby pixels within a 20kb radius. If found, these pixels were grouped into a loop cluster, and their centroid was calculated. This process was repeated iteratively until all loops were processed. For each cluster, the pixel with the lowest q-value was designated as the cluster summit, representing the final non-redundant loop for that resolution. This step was performed for both 10kb and 25kb identified loops. Then, the non-redundant loop lists from the 10kb and 25 kb resolutions were then merged. During this step, if a 25kb loop cluster overlapped with a 10kb loop cluster, the 25kb loop was removed to avoid redundancy and prioritize higher-resolution data.

#### Loop partition

Structural loops were defined as those with both anchors occupied by CTCF and cohesin ChIP-seq peaks and at most 1 anchor occupied by CREs.

Structural loops were classified hierarchically based on their nesting relationships. A singleton loop is a structural loop that exists independently, neither containing nor being contained within another loop. A loop that is nested within another but contains no loops of its own is classified as a Tier 1 primary structural loop. A Tier 2 structural loop directly encompasses a Tier 1 loop without any intermediate loops. This classification scheme extends to higher tiers, where each tier represents a structural loop that directly contains a loop from the tier immediately below it.

Primary tier1 structural loops were further classified based on their local chromatin contexts. For this analysis, the internal ChIP-seq signals for H3K27ac, H3K36me3, H3K27me3, and H3K9me3 were quantified for each structural loop. Each histone modification signal was normalized genome-wide, transformed into z-scores, and then subjected to unsupervised k-means clustering. This clustering resulted in three distinct clusters:

Cluster 1: Characterized by enrichment of H3K27ac, indicative of active euchromatin. Cluster 2: Enriched for H3K27me3, corresponding to facultative heterochromatin.

Cluster 3: Marked by H3K9me3 enrichment, representing constitutive heterochromatin.

#### Aggregated plots for loops and domains

Aggregated plots were generated using the Python package Coolpup (v1.1.1). For unscaled aggregated peak analysis, loops smaller than 20kb were removed from the plots to avoid influence from pixels close to the diagonal.

#### TT-seq preprocessing and differential expression analysis

TT-seq reads were processed to quantify gene expression in a strand-specific manner. First, sequencing reads were aligned to the longest RefSeq-annotated mm9 mouse reference genome transcript set using STAR v2.7.10b. After alignment, quality control and filtering were performed with Samtools v1.6 using the parameters “-q 10 -f 0x2” to remove low-quality alignments. PCR duplicates were subsequently removed using Picard v2.25.5. Next, the strand-specific BAM files were separated into forward and reverse strands. Forward-strand reads were isolated using the Samtools parameters “-f 128 -F 16 -f 80”, and reverse-strand reads were isolated using “-f 144 -f 64 -F 16”. BigWig files were then generated using the bamCoverage function in deepTools v3.5.4. The multicov function in bedtools v2.31.1 was used to obtain fragment counts across genomic features, which served as the basis for gene quantification. For positive-strand genes, forward-strand reads were quantified, while for negative-strand genes, reverse-strand reads were used.

Differential expression analysis was carried out in R using the DESeq2 package v1.42.1. Genes were classified based on their statistical significance and fold change. Specifically, genes with an adjusted P value (padj) < 0.05 and a log_2_ fold change (log_2_FC) > log_2_(1.5) were identified as upregulated. Conversely, genes with a padj < 0.05 and a log_2_FC < -log_2_(1.5) were considered downregulated. Genes that did not meet these criteria were classified as non-differentially expressed.

#### ABC model predicts enhancer-promoter interactions

To predict enhancers of active genes and establish promoter–enhancer (P–E) and promoter–promoter (P–P) connections, we applied the Activity-by-Contact (ABC) model (https://github.com/broadinstitute/ABC-Enhancer-Gene-Prediction)^52^.Enhancer activity was estimated using H3K27ac ChIP-seq and ATAC-seq signals from asynchronously growing G1E-ER4 cells. This dataset was combined with replicate-merged Hi-C datasets from *Stag1*^mAID^, *Stag2*^mAID^, and *Stag1*^mAID^*Stag2*^mAID^ cells. For each sample, a 5kb resolution Hi-C contact matrix was used to calculate the contact frequency between each regulatory element and its target gene.

Enhancer activity was assessed across two post-mitotic cell-cycle stages, with or without 5-Ph-IAA treatment: early G1 (“–5-Ph-IAA” and “+5-Ph-IAA”) and late G1 (“–5-Ph-IAA” and “+5-Ph-IAA”), to predict P–E and P–P connections between enhancers and their target genes. The ABC score for an element on a gene was calculated by multiplying the enhancer activity by the contact frequency between the enhancer and the gene, and dividing this product by the sum of such products for all candidate elements within a 5Mb window. Increasing the ABC score threshold reduces the number of interactions classified as high-confidence.

An ABC score threshold of 0.025 was applied consistently across all samples, and predicted connections from each sample were combined to generate a set of high-confidence P–E and P–P connections. An interaction was considered valid if its ABC score met the threshold in at least one sample. Each P–E or P–P pair was then mapped to its target gene and classified based on differential expression status (upregulated, downregulated, or non-significant).

#### Calculation of intra-loop contact frequency

To determine the intra-loop contact frequency, pixels within each structural loop domain were extracted. The average observed/expected value across all pixels within a given loop domain was then calculated. This average represents the intra-loop contact frequency for that specific loop.

#### Data availability

The Hi-C and ChIP-seq data generated in this study will be deposited in the GEO database under the accession number GSE306128. External ChIP-seq datasets used in this study include: H3K27ac (GSE61349)^53^, H3K36me3 (GSM946529), H3K9me3 (GSM946542), H3K27me3 (GSM946531)^54^, CTCF (GSE129997), and Rad21 (GSE129997)^24^. External Hi-C datasets include late-G1 phase Nipbl-depleted cells (GSE256073)^38^ and mid-G1 phase CTCF-depleted cells (GSE168251)^33^.

## Code availability

Codes used in this study are available at Zenodo: 10.5281/zenodo.10968270 (sequencing analysis pipelines).

## Figure Legends

**Extended Data Figure 1:**
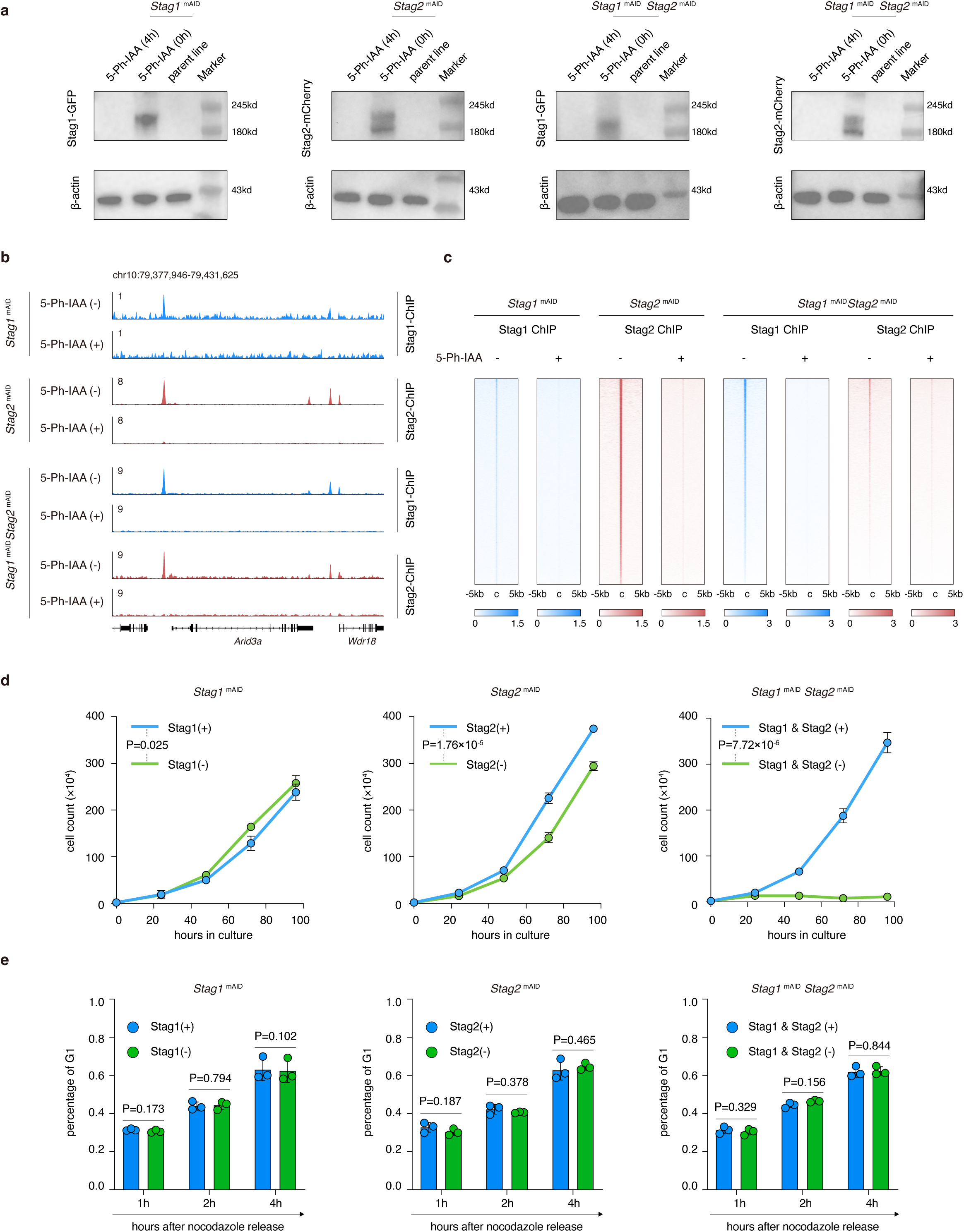
Characterization of *Stag1*ᵐᴬᴵᴰ, *Stag2*ᵐᴬᴵᴰ, and *Stag1*ᵐᴬᴵᴰ *Stag2*ᵐᴬᴵᴰ cell lines. **a**, Western blot analysis of *Stag1*ᵐᴬᴵᴰ, *Stag2*ᵐᴬᴵᴰ, and *Stag1*ᵐᴬᴵᴰ *Stag2*ᵐᴬᴵᴰ cells treated with or without 5-Ph-IAA. Anti-GFP and anti-mCherry antibodies were used to detect Stag1 and Stag2 respectively. **b**, Genome browser tracks showing loss of Stag1 or/and Stag2 peaks upon 5-Ph-IAA treatment in *Stag1*ᵐᴬᴵᴰ, *Stag2*ᵐᴬᴵᴰ, and *Stag1*ᵐᴬᴵᴰ *Stag2*ᵐᴬᴵᴰ cells respectively. **c**, Metaplots of Stag1 or Stag2 ChIP-seq signals centered at peak summits (±5 kb) for the indicated cell lines with or without 5-Ph-IAA treatment. **d**, Growth curves showing the impacts of Stag1 or/and Stag2 depletion on cell proliferation. Data are mean ± S.D. from three independent biological replicates; *P* values were calculated using two-way repeated measures ANOVA with Tukey’s multiple comparisons test. **e**, Bar graphs showing the percentage of G1 phase cells after nocodazole release for indicated durations after Stag1 or/and Stag2 depletion. Data are mean ± S.D. from three biological replicates; *P* values are calculated using student’s t-test.

**Extended Data Figure 2:**
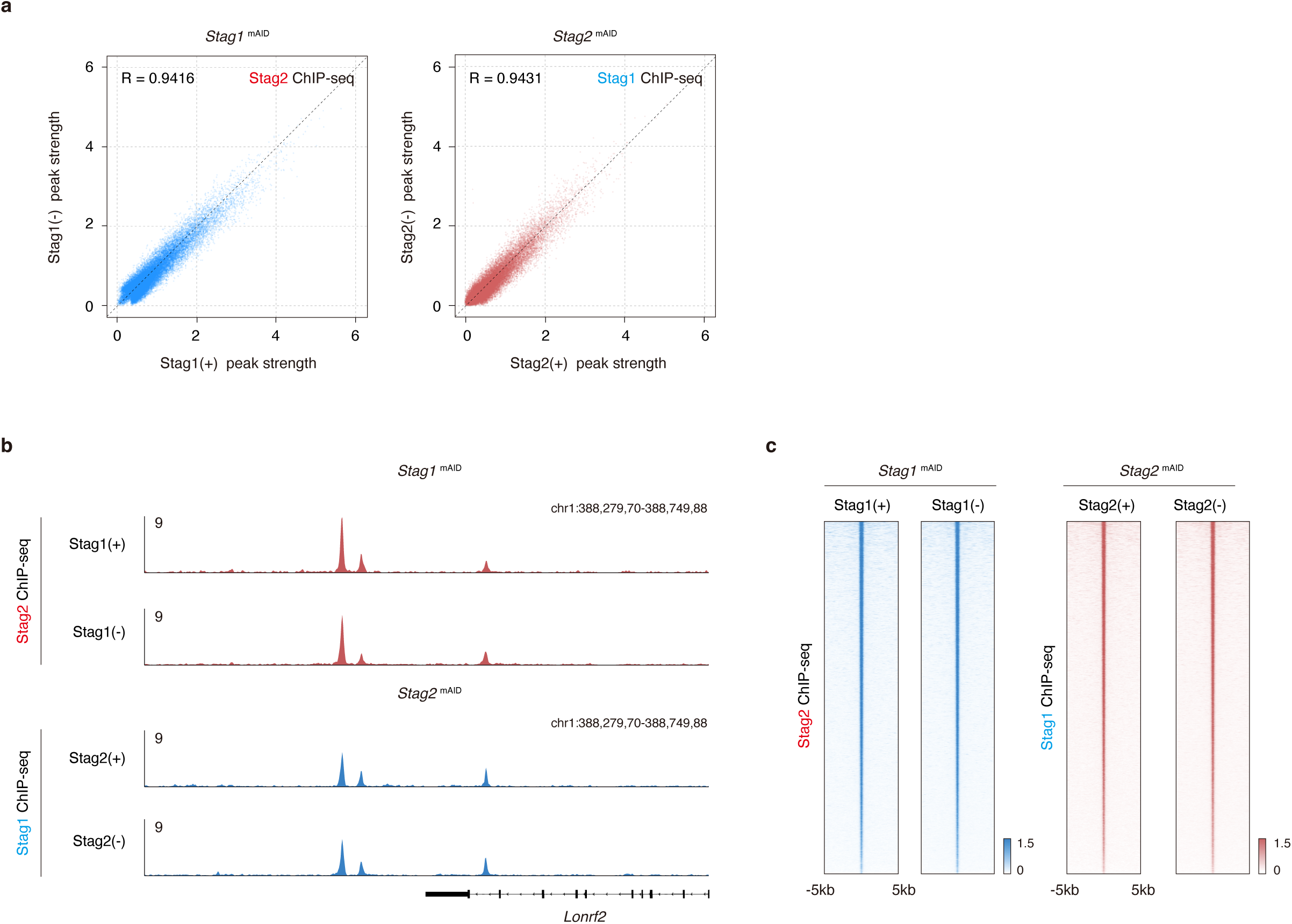
Stag1 and Stag2 chromatin binding profiles are independent of each other. **a**, Scatter plots showing that Stag2 ChIP-seq peak strength is not measurably perturbed upon Stag1 depletion (Left panel) and vice versa (Right panel). **b**, Representative genome browser tracks showing that Stag2 ChIP-seq binding profile is not changed upon Stag1 depletion (Upper panel) and vice versa (Lower panel). **c**, Metaplots showing that Stag2 ChIP-seq peak strength is not measurably perturbed upon Stag1 depletion (Left panel) and vice versa (Right panel).

**Extended Data Fig. 3:**
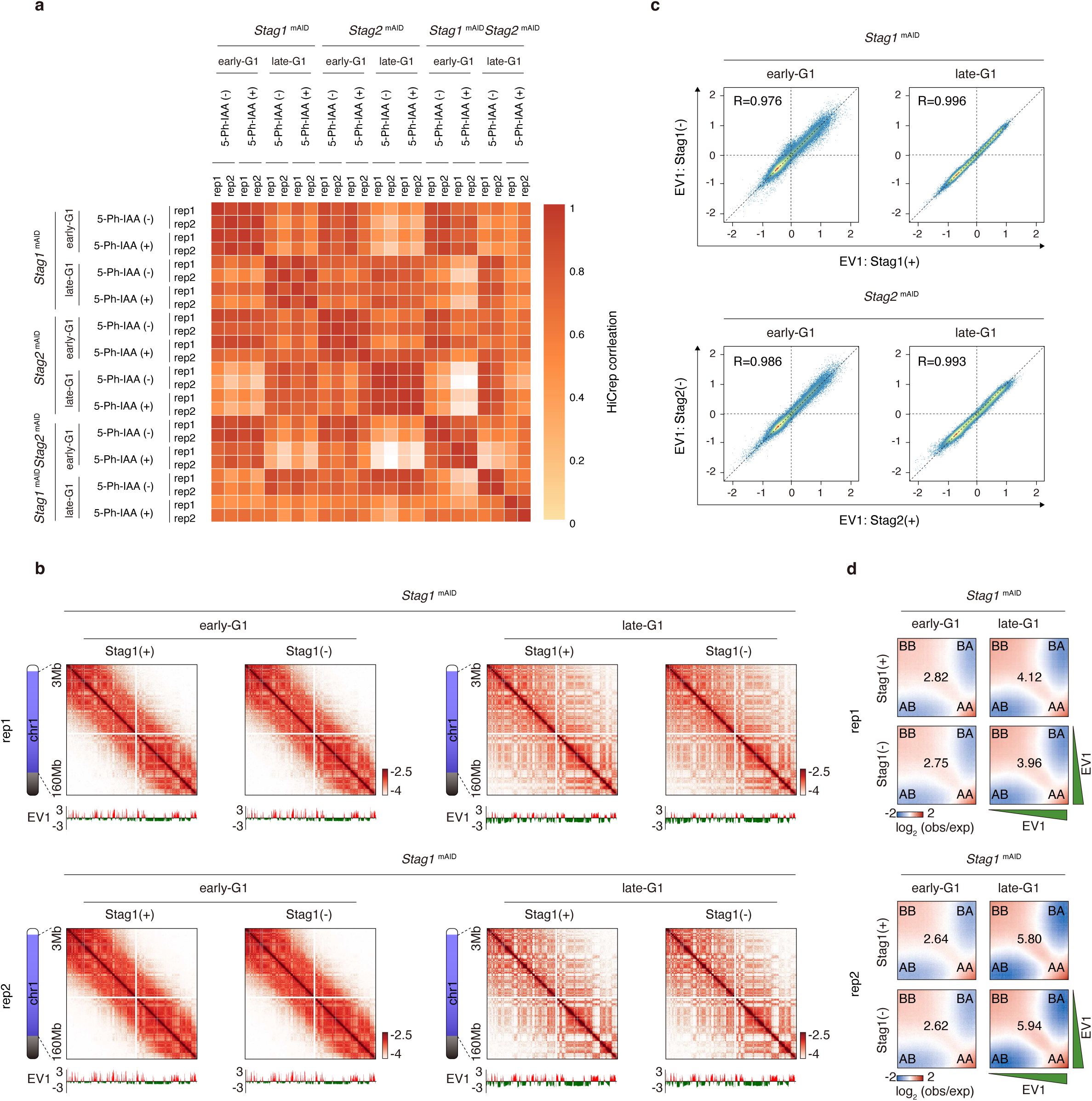
Hi-C reproducibility and compartment analysis of Stag1- or Stag2-deficient cells after mitosis. **a**, HiCRep results showing the high reproducibility of Hi-C replicates for each condition. **b**, Representative Hi-C contact maps of individual biological replicates showing the post-mitotic reformation of compartments upon Stag1 depletion. **c**, Scatter plots showing highly correlated EV1 values with or without Stag1 or Stag2 in early- and late-G1 phase. **d**, Saddle plots of individual biological replicates with or without Stag1.

**Extended Data Figure 4:**
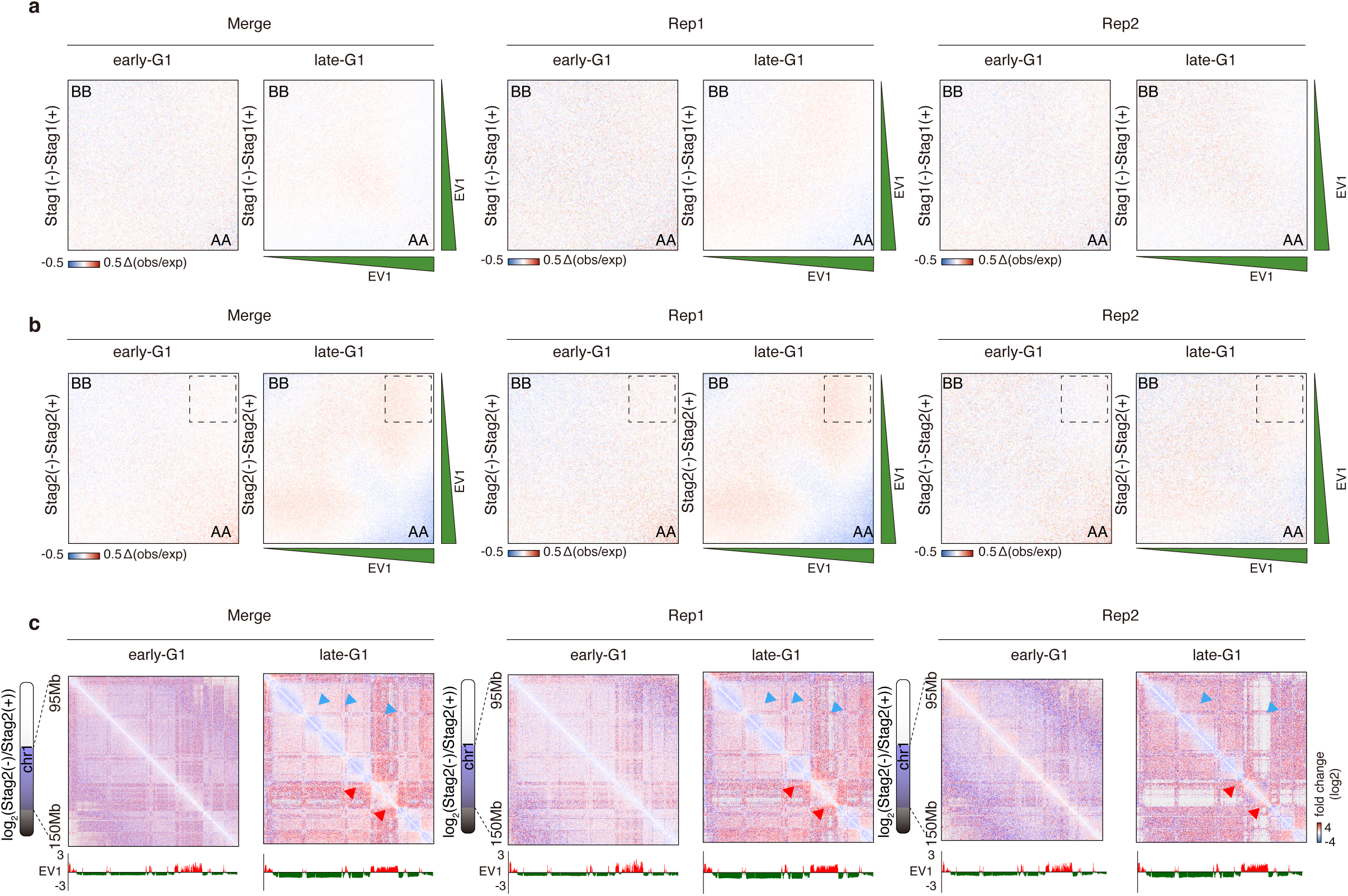
Stag2 depletion mildly affects chromatin re-compartmentalization after mitosis. **a**, Subtracted Saddle plots of Stag1-depleted and replete samples, showing minimally affected chromatin compartments upon Stag1 depletion in early- and late-G1 phase. **b**, Subtracted Saddle plots of Stag2-depleted and replete samples, showing a slight reduction in A/B segregation in late-but not early-G1 phase upon Stag2 loss. **c**, Representative Hi-C contact maps showing the changes in chromatin-compartmentalization upon Stag2 loss in early- and late-G1 phase. Gain of B-B compartmental interactions and loss of A-B segregation are indicated by blue and red arrows respectively. Results for replicate-merged and individual biological replicate samples were shown.

**Extended Data Figure 5:**
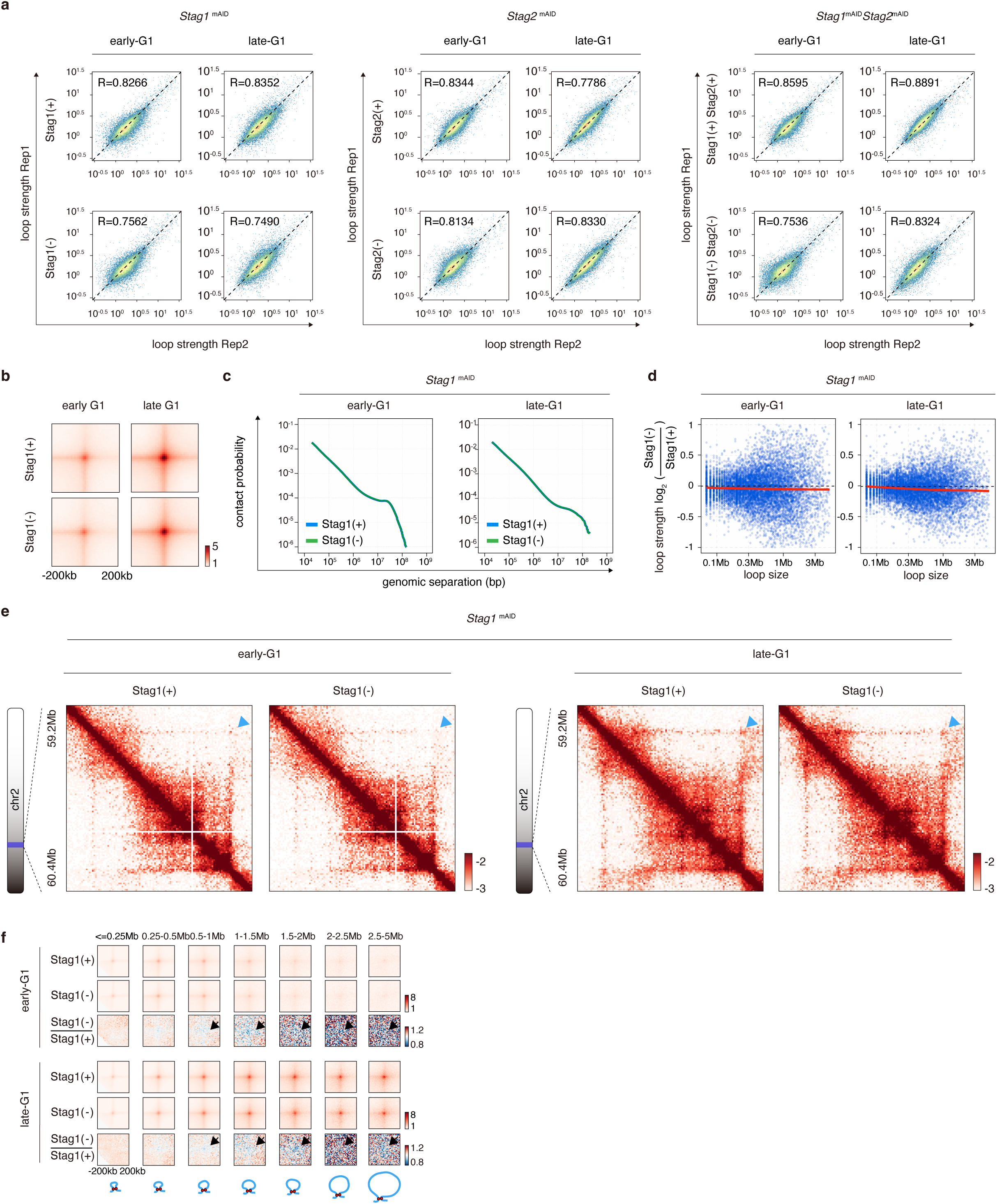
Stag1 depletion mildly reduces post-mitotic chromatin loop reformation. **a**, Scatter plots showing the correlation of structural loop strength between biological replicates. **b**, APA plots showing that structural loops are unaffected by Stag1 depletion in early- and late-G1 phase. **c**, *P(s)* curve is not changed by Stag1 depletion in early- and late-G1 phase. **d**, Scatter plot showing the highly comparable structural loop strength in early- and late-G1 phase cells with or without Stag1. **e**, KR-balanced Hi-C contact maps showing that structural loop signal is marginally reduced by Stag1 depletion in early- and late-G1 phase (cyan arrows). **f**, APA plots showing that Stag1 depletion mildly reduces structural loops, with a slight reduction observed in larger loops (black arrows), in early-G1 and late-G1 phases.

**Extended Data Figure 6:**
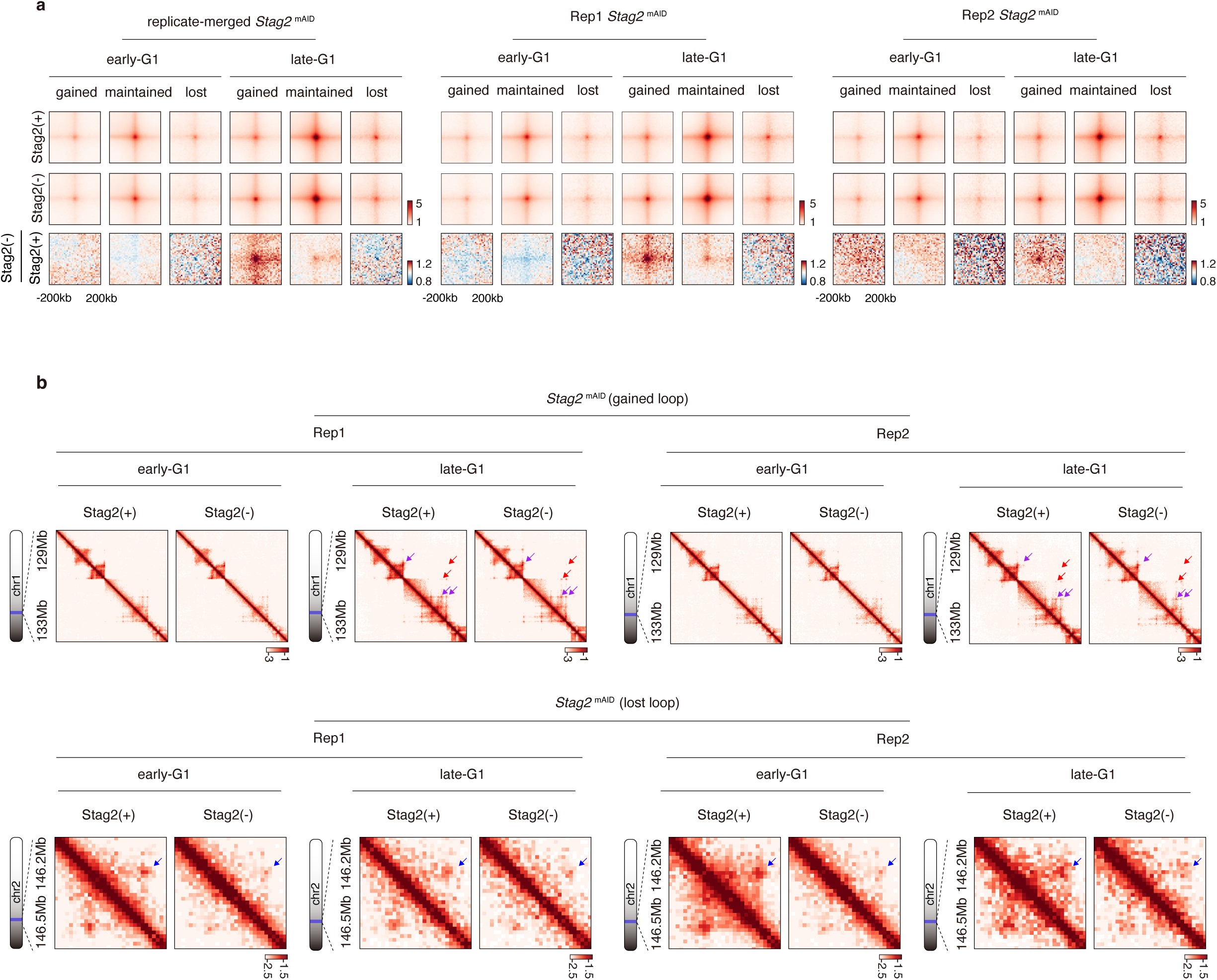
Stag2 depletion affects post-mitotic structural loop reformation. **a**, APA plots showing the changes of lost, maintained and gained loops upon Stag2 depletion in early- and late-G1 phase cells. Results of replicate-merged and individual replicates are shown. **b**, Hi-C contact maps showing representative gained and lost loops upon Stag2 depletion in early- and late-G1 phase. lost, maintained and gained loops are indicated by blue, purple and red arrows respectively. Results of individual replicates are shown.

**Extended Data Figure 7:**
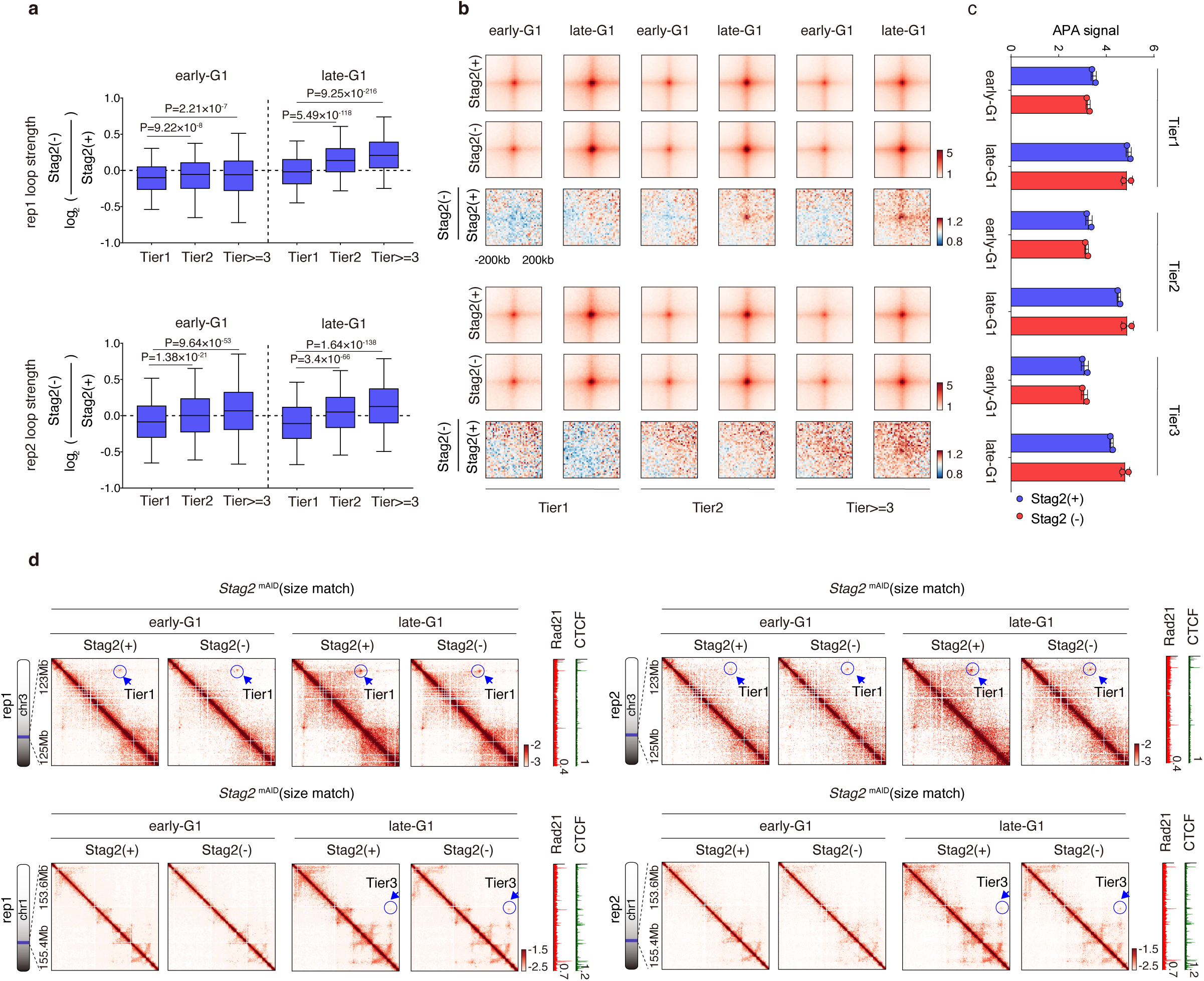
Impacts of Stag2 loss on structural loops of different hierarchical tiers. **a**, Boxplots showing the impacts of Stag2 loss on loops across different tiers. Log_2_FC values are calculated for each biological replicate respectively. Data for early- and late-G1 phase are both shown. *P* values are calculated using a two-sided Wilcoxon rank-sum test. **b**, APA plots showing the changes in structural loops across different tiers upon Stag2 loss in early- and late-G1 phase. Data for two biological replicates are shown independently. **c**, Bar graphs showing the quantification of APA plots in (**b**). **d**, KR-balanced Hi-C contact maps showing representative size-matched Tier1 and Tier3 structural loops in early- and late-G1 phase cells with or without Stag2. Maps for individual biological replicates are shown. Late-G1 Rad21 and CTCF ChIP-seq tracks are shown on the right.

**Extended Data Figure 8:**
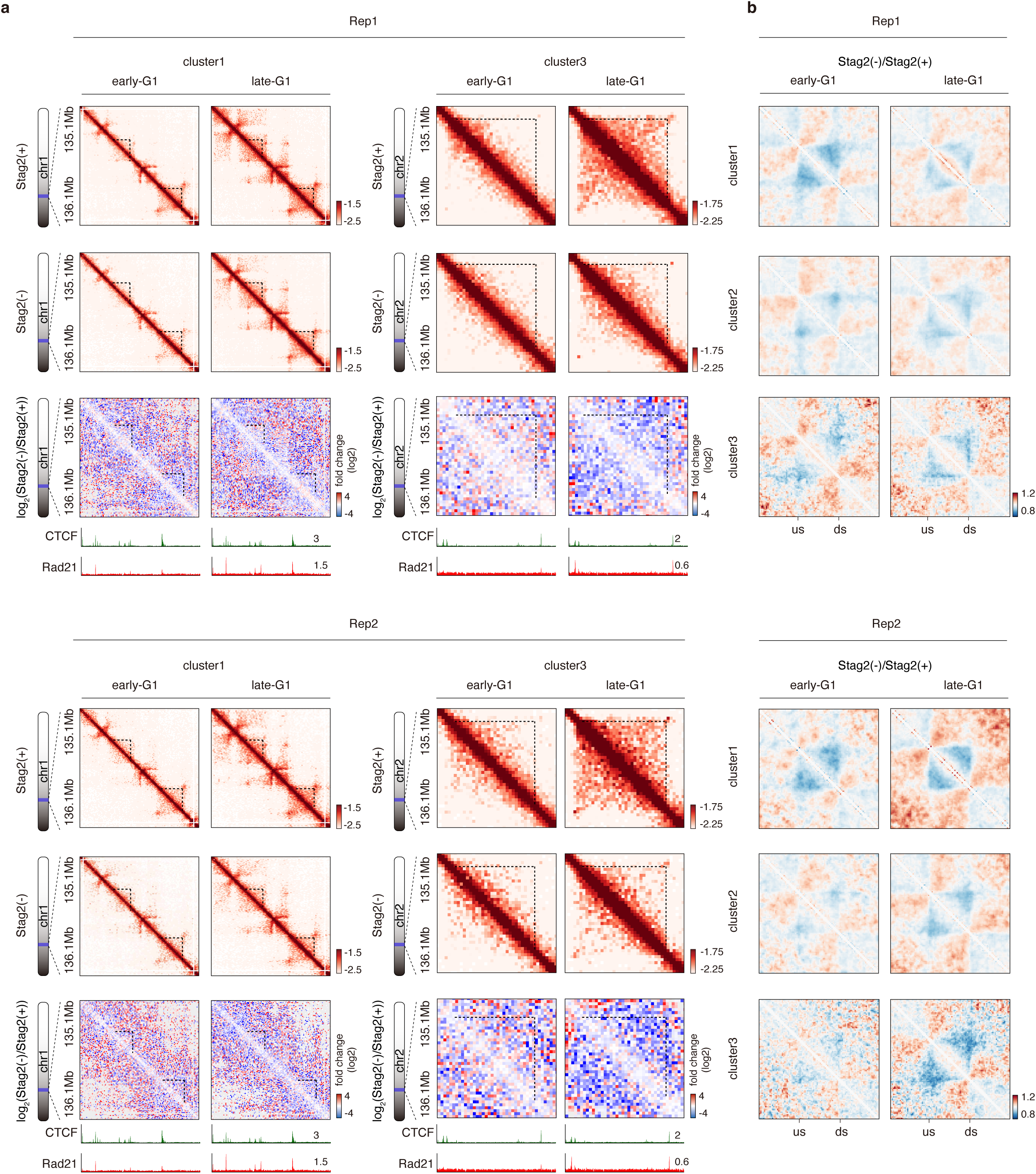
Additional data supporting chromatin context-dependent regulation of loop extrusion by Stag2. **a**, KR-balanced Hi-C contact maps showing the changes in cluster 1 (Left panel) and cluster 3 (right panel) loops upon Stag2 depletion in early- and late-G1 phase. Dotted lines demarcate intra-loop contacts. Data for individual biological replicates are shown. Contact maps are coupled with corresponding ChIP-seq tracks for CTCF and Rad21. **b**, Compiled and rescaled contact maps showing the distinct impact of Stag2 loss on the intra-loop contacts across all three clusters. “us” and “ds” denotes the upstream and downstream loop anchors, respectively. Plots for individual biological replicates are shown.

**Extended Data Figure 9:**
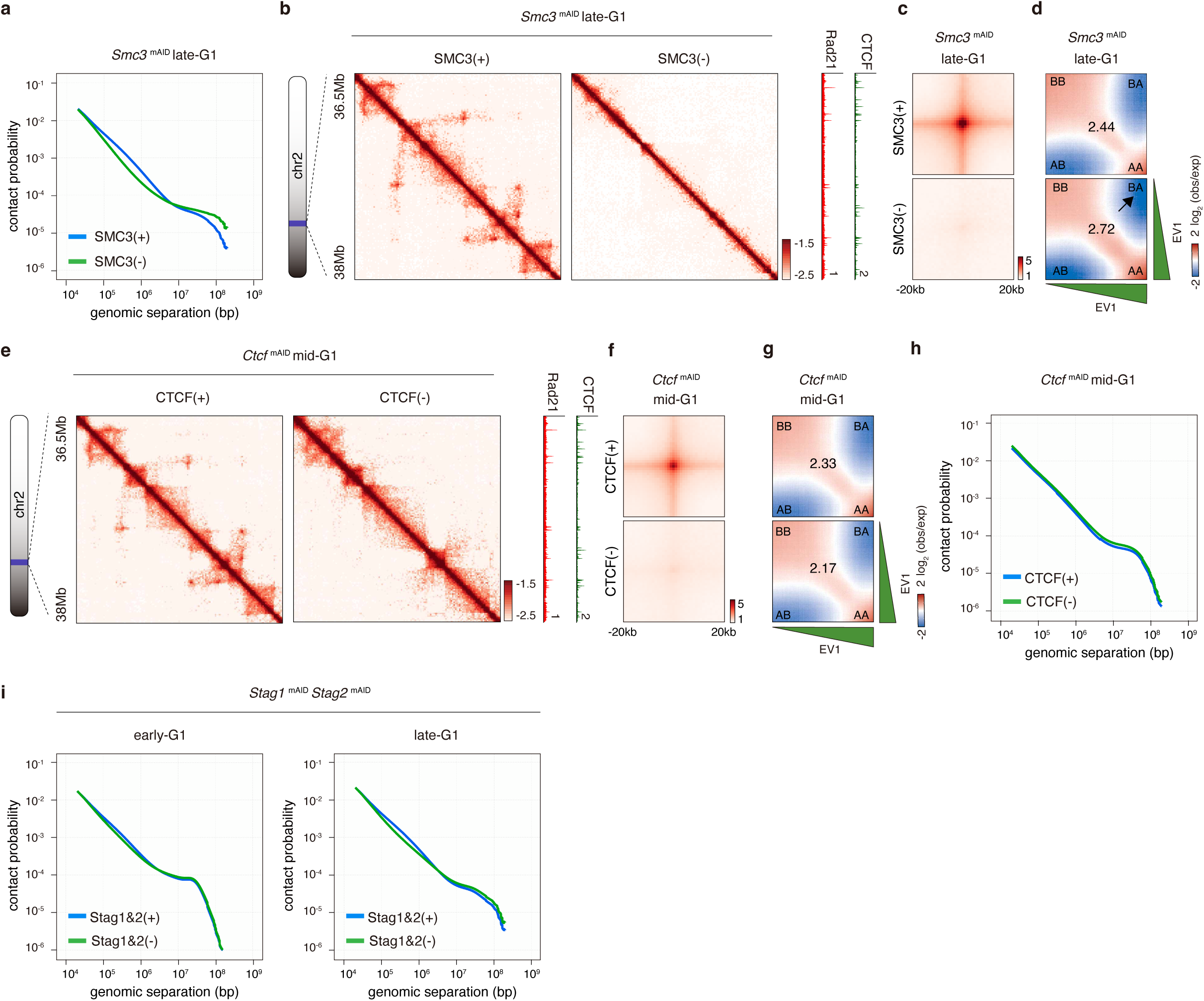
Effects of SMC3, CTCF, and Stag1/2 depletion on chromatin architecture after mitosis. **a**, *P(s)* curves showing dramatically reduced short-range contacts (loop and TAD scale) upon SMC3 depletion, confirming loss of cohesin loop extrusion. **b**, KR-balanced Hi-C contact maps showing the complete loss of structural loops in the late-G1 phase upon SMC3 depletion. The Hi-C contact maps are coupled with late-G1 phase CTCF and Rad21 ChIP-seq tracks. **c**, APA plots showing the complete loss of structural loops in the late-G1 phase upon SMC3 depletion. **d**, Saddle plots showing that chromatin compartmentalization is strengthened upon SMC3 depletion. Black arrow indicates enhanced A/B separation upon SMC3 loss. **e**, KR-balanced Hi-C contact maps in mid-G1 phase showing that structural loops are lost upon CTCF depletion. **f**, APA plots showing the loss of structural loops upon CTCF depletion in mid-G1 phase. **g**, Saddle plots showing that chromatin compartmentalization is not affected in CTCF-depleted cells in mid-G1 phase. **h**, *P(s)* curves of CTCF-replete and depleted samples in mid-G1 phases. **i**, *P(s)* curves showing reduced short-range contacts (loop and TAD scale) upon Stag1 and Stag2 co-depletion in both early-G1 and late-G1 phases.

**Extended Data Figure 10:**
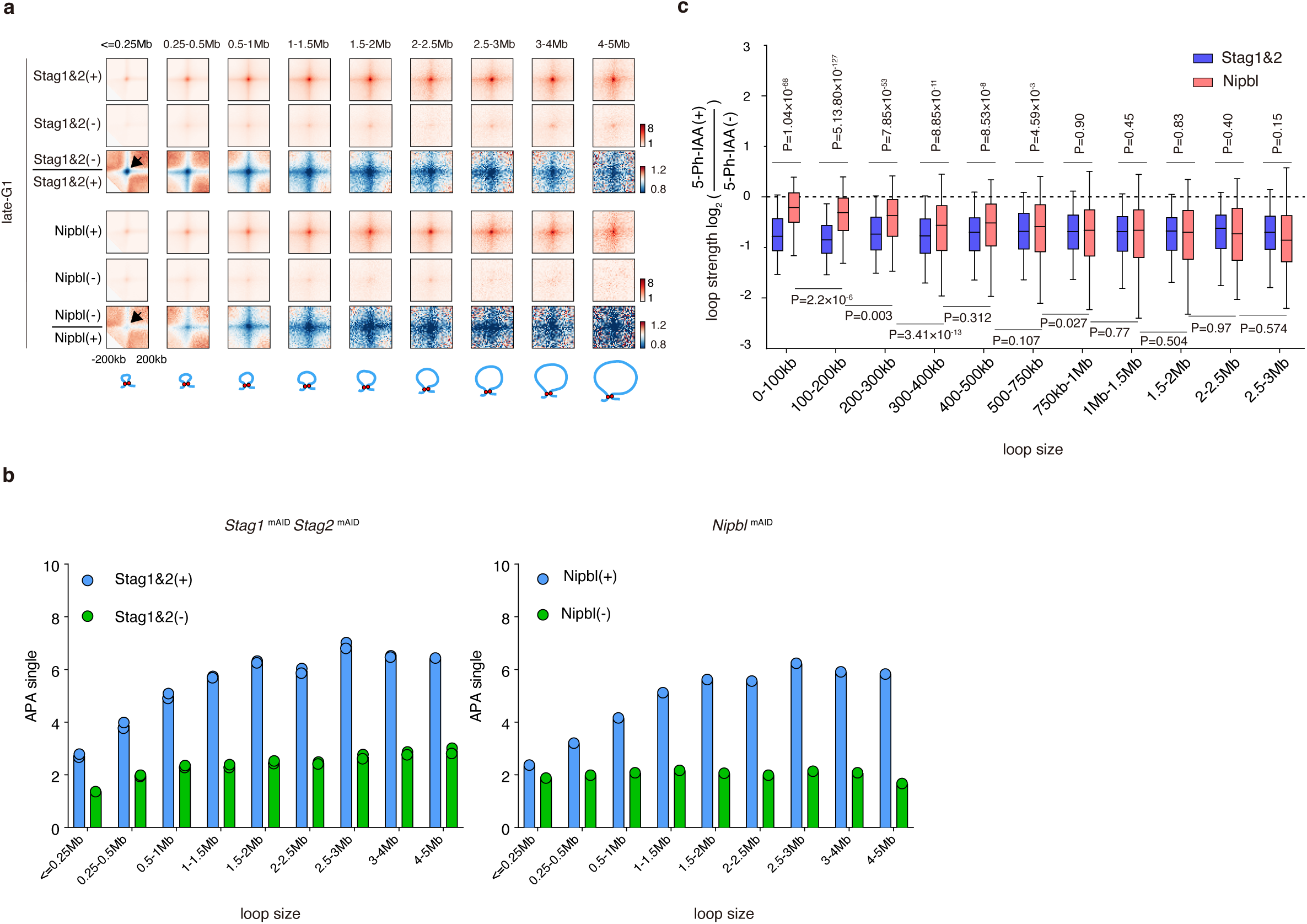
Stag1 and Stag2 co-depletion does not affect the extruding processivity of chromatin-associated cohesin. **a**, APA plots, showing that Stag1 and Stag2 co-depletion reduces structural loops uniformly, regardless of their size (black arrow, upper panel). In contrast, Nipbl depletion, which affects cohesin extrusion capacity, causes a size-dependent loss of loops, with smaller loops being less affected (black arrow, lower panel). **b**, Bar plots showing the quantification of APA plots in (**a**). For Stag1 and Stag2 co-depletion, each dot represents a biological replicate. For Nipbl depletion, data are shown for the replicate-merged samples. **c**, Boxplots, showing the log_2_FC in structural loop strength, revealing the distinct effects of Stag1 and Stag2 co-depletion versus Nipbl depletion on loops of various sizes. *P* values are calculated using two-sided Wilcoxon rank-sum test. Note that paired comparison was used when comparing the same set of loops across two different conditions.

**Extended Data Figure 11:**
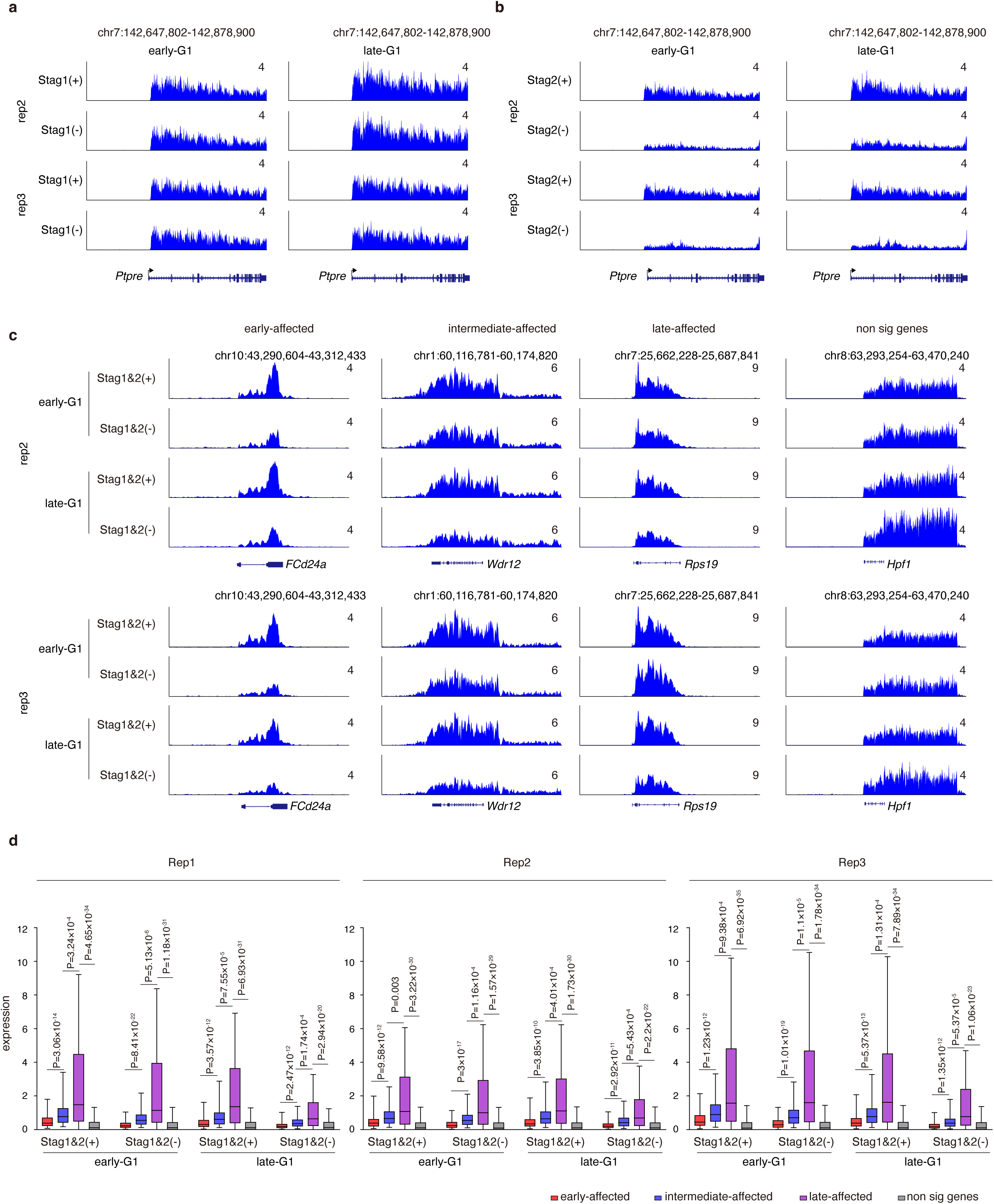
Effects of Stag1 or/and Stag2 loss on post-mitotic transcription reactivation. **a,** Genome browser tracks showing the TT-seq signals of additional biological replicates at the *Ptpre* locus in early- and late-G1 phase cells with or without Stag1. **b**, Similar to (**a**) showing the TT-seq signals of additional biological replicates at the *Ptpre* locus in early- and late-G1 phase cells with or without Stag2. **c,** Genome browser tracks showing TT-seq signals of additional biological replicates upon Stag1 and Stag2 co-depletion. Tracks of four representative loci: *FCd24a* (early-affected), *Wdr12* (intermediate-affected), *Rps19* (late-affected), and *Hprt1* (non-significant) were shown. **d,** Boxplots showing TT-seq signals in the gene body for early-affected, intermediate-affected, late-affected and non-significant gene groups. Data are shown for early- and late-G1 cells, with or without Stag1 and Stag2. Results for three independent biological replicates were shown. *P* values were calculated using two-sided Wilcoxon rank-sum test.

## Supplementary Tables

Supplementary Table 1: Read statistics of Hi-C experiments

Supplementary Table 2: ligo lists

